# High- and Low-Frequency Deep Brain Stimulation in the Subthalamic Nucleus differentially modulate Response Inhibition and Action Selection in Parkinson’s Disease

**DOI:** 10.1101/2022.05.13.491771

**Authors:** Josefine Waldthaler, Alexander Sperlich, Aylin König, Charlotte Stüssel, Frank Bremmer, Lars Timmermann, David Pedrosa

## Abstract

**Background:** While deep brain stimulation (DBS) in the subthalamic nucleus (STN) improves motor functions in Parkinson’s disease (PD), it has also been associated with increased impulsivity.

**Methods:** A combined approach of eye-tracking and high-density EEG was used to investigate how high- and low-frequency DBS impact impulsive actions in the antisaccade task in a cohort of ten persons with PD. Computational modelling of the behavioral outcomes allowed a nuanced insight into the effect of DBS on response inhibition and action selection processes. Results: Against our expectations, both 130 Hz- and 60 Hz-DBS improved response inhibition as both resulted in a reduced rate of early reflexive errors. Correspondingly, DBS with both frequencies led to increased desynchronization of beta power during the preparatory period which may be a correlate of anticipatory activation in the oculomotor network.

Low-frequency DBS additionally was associated with increased midfrontal theta power, an established marker of cognitive control. While higher midfrontal theta power predicted longer antisaccade latencies in off-DBS state on a trial-by-trial basis, 130 Hz-DBS reversed this relationship. As informed by the computational model, 130 Hz-DBS further led to a shift in the speed-accuracy trade-off causing an acceleration and error-proneness of actions later in the trial.

**Conclusions:** Our results disentangle the impact of DBS on early and late impulsive actions. Only 130 Hz-DBS may disrupt theta-mediated cognitive control mechanisms via medial frontal – STN pathways that are involved in delaying action selection. 60 Hz-DBS may provide beneficial effects on response inhibition without the detrimental effect on action selection seen with 130 Hz-DBS.

**Funding:** This study was supported by the SUCCESS program of Philipps-University Marburg (JW), the Hessian Ministry of Sciences and the Arts, clusterproject: The Adaptive Mind – TAM (FB / AK) and the German Research Foundation (DFG). International Research Training Group 1901 (FB / AK)

## INTRODUCTION

The term performance monitoring defines a set of several cognitive functions that underlie the ability to adjust behaviors according to internal or environmental demands (Ullsperger, 2006). Response inhibition, i.e., withholding prepotent reflexive responses and thereby allocating more time to shape the behavioral strategy according to context often results in more favorable outcomes of our actions (Obeso et al., 2011). Impaired response inhibition, on the other side, leads to impulsivity, i.e., the tendency of acting without delay, reflection or voluntary directing (Bari & Robbins, 2013).

Since response inhibition is modulated by activation of dopamine-dependent fronto-striatal networks, these cognitive functions are particularly affected by an aberrant dopaminergic system in Parkinson’s disease (PD) (Kudlicka et al., 2011). In fact, impaired executive functioning as well as impulsive and compulsive behaviors (ICB) are commonly encountered in PD affecting approximately 40 %, respectively 14% of patients (Godefroy et al., 2010; Weintraub et al., 2010). Impulsivity, and in particular behavioral addictions, so called Impulse Control Disorders (ICD), negatively impact health and quality of life of affected patient and their relatives likewise (Voon et al., 2017).

Deep Brain Stimulation (DBS) of the subthalamic nucleus (STN) is an effective treatment in PD. While STN-DBS improves motor symptoms, a variety of behavioral studies have lend credence to stimulation-induced impaired response inhibition (Ballanger et al., 2009; Hershey et al., 2004; Obeso et al., 2013; N. J. Ray et al., 2009; Witt et al., 2004). In this study, we aim at exploring the effects of different DBS pulse frequencies with respect to switched-off stimulation on the antisaccade task, an established paradigm assessing response inhibition.

### Neural correlates of response inhibition

Studies on healthy participants revealed that motor response inhibition activates a network consisting of prefrontal and premotor regions along with the basal ganglia (Stevens et al., 2007), all of which interact via frequency-specific synchronized neuronal oscillations. In brief, main cortical areas involved in the response inhibition network are the inferior frontal gyrus (iFG), the medial and anterior cingulate cortex (ACC), and the dorsolateral prefrontal cortex (DLPFC). Here, ACC seems to be determinant for delaying responses whenever conflicts occur or in cases of demand for cognitive control as it allows more time for successful action selection (Hinault et al., 2019). In other words, the ACC serves a *proactive* response inhibition. The iFG, on the contrary, is particularly involved in general stopping of ongoing activity after actions have already been selected (Chikazoe et al., 2007), i.e. *reactive* response inhibition. However, these two mechanisms may share parts of a common network structure (Zhang & Iwaki, 2019) and work in parallel, as proposed by Wiecki and Frank’s model of inhibitory control (Wiecki & Frank, 2013).

Communication between frontal brain areas occurs via theta oscillations (4-8 Hz) over medial frontal regions (referred hereafter as midfrontal theta), which are detectable in tasks requiring cognitive control and response inhibition (cf. Cavanagh & Frank, 2014, for review). Thus, midfrontal theta has been proposed as neural signature of an action monitoring system of the brain. Midfrontal theta is generated by ACC and the pre-supplemental motor area (pre-SMA) as intracranial recordings in non-human primates and humans as well as fMRI studies suggest (Cohen et al., 2008; Hauser et al., 2014; Tsujimoto et al., 2006; Wang et al., 2005). In PD, midfrontal theta activity is diminished during cognitive control (Singh et al., 2018).

Yet, it is undisputable that assigning cognitive control merely to cortical areas would pose an oversimplification. On a subcortical level, the basal ganglia are critically involved in the process of response inhibition serving as a system for response selection and initiation. In particular, and in accordance with classical models of cortico-basal ganglia circuitry, activity in the indirect pathway via STN inhibits prepotent responses to external cues until the selected response is triggered via the direct pathway (Chevalier & Deniau, 1990; Redgrave et al., 1999). These dynamic properties of the STN to delay action selection when accuracy is favored over speed or when actions need to be selected out of more than one simultaneously activated response sets are pivotal for efficient and successful response inhibition (Frank, 2006; Herz et al., 2018). Frequency-specific STN activity seems to play a role in both reactive inhibition as well as in the implementation of the proactive “hold your horses” signal (Benis et al., 2014). Studies with parallel local field potential (LFP) and EEG recordings suggested that these processes are associated with changes of beta band activity (13-30 Hz) in and synchronization between the iFG and the STN (Alegre et al., 2013; Schaum et al., 2021; Swann et al., 2011), respectively theta band activity in and synchrony between the medial frontal cortex and STN (Ray et al., 2012; Zavala et al., 2013, 2016) (cf. B. Zavala et al., 2015, for review).

### Effects of Deep Brain Stimulation on response inhibition

Growing evidence indicates that chronic subthalamic DBS may interfere with different aspects of impulsivity in PD. Whereas initial findings indicated detrimental effects on response inhibition with high-frequency pulses (Hershey et al., 2004; Jahanshahi et al., 2000), subsequent larger studies reported no effect, or even improved inhibitory control under STN-DBS in PD. A review by Scherrer et al. gathered sufficient evidence to support the notion of STN-DBS increasing impulsivity under speed pressure or at conflict, i.e., when decision-making between competing choices is required (Scherrer et al., 2020).

The mechanisms underlying this disinhibition remain speculative. According to fMRI studies, activation of DBS is associated with impairing response inhibition and simultaneously reducing activity in dorsal ACC, iFG and pre-SMA (Ballanger et al., 2009) corroborating the aforementioned theoretical framework. EEG studies evaluating effects of STN-DBS on cognitive control, which may overcome the low temporal discrimination of fMRI, are technically difficult. Thus, results are sparse and inconclusive. For example, beta band power over the right prefrontal cortex was higher during a stop signal task with DBS switched on compared to off-DBS state which was accompanied by *improved* response inhibition (Swann et al., 2011). Another recent EEG study found neither effects of STN-DBS on proactive response inhibition nor on the activity of the underlying cortical network compared to a non-operated PD group (De Pretto et al., 2021).

### The Antisaccade task

The antisaccade task is an established eye-tracking paradigm to explore response inhibition. Participants are asked to inhibit a reflexive saccade in the direction of a visual stimulus (the *pro*saccade) and to execute a voluntary saccade in the opposite direction instead (the *anti*saccade) (Figure 1A). Both pro- and antisaccades activate a well-known widespread network associated with planning and execution of saccades, encompassing frontal, parietal and supplementary eye fields (FEF, PEF, respectively SEF), thalamus, striatum, and the intraparietal cortex (Jamadar et al., 2013). Antisaccades additionally recruit the right DLPFC, and ACC (Brown et al., 2007) which seem to be essential to successfully suppress reflexive prosaccade errors top-down (Pa et al., 2014). Moreover, preparation for successful antisaccades induces an increase in midfrontal theta power and fronto-central inter-trial theta coherence compared with both no-go trials and errors (Cordones et al., 2013; van Noordt et al., 2017) further supporting the idea of increased top-down cognitive control.

**FIGURE 1:**
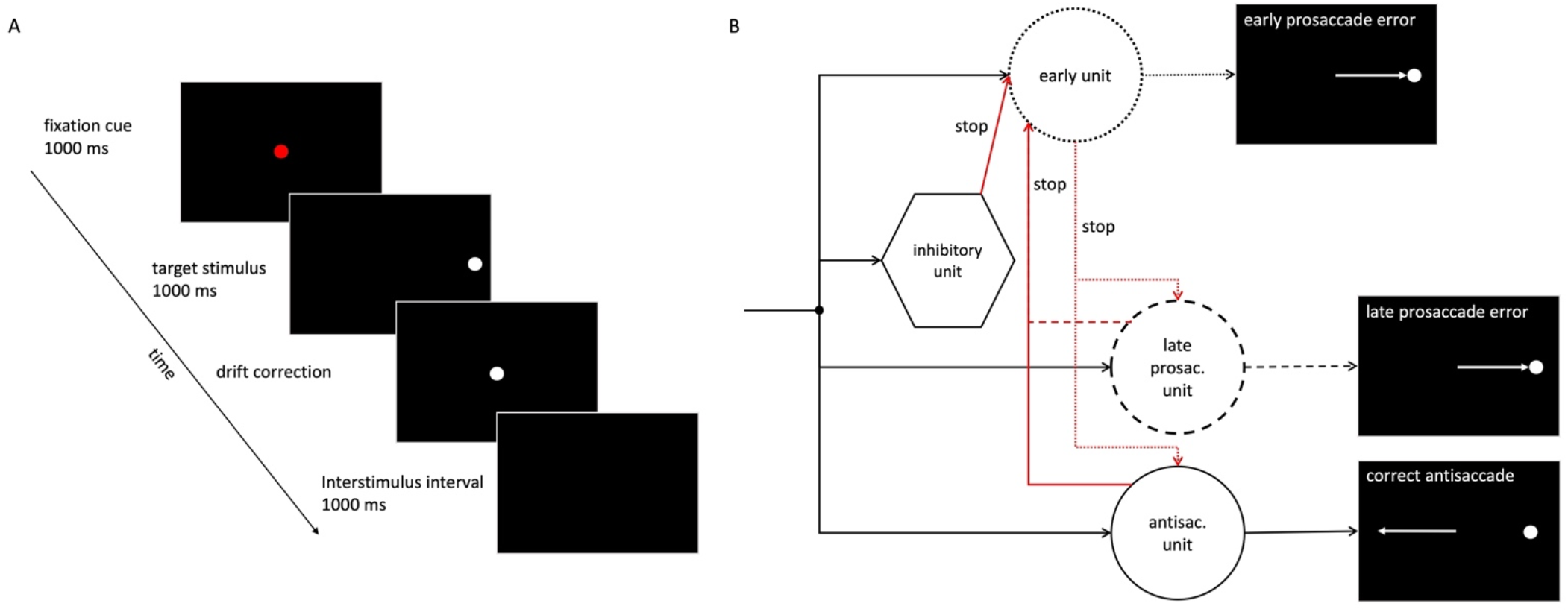
1A: The antisaccade task. Please see section Eye-tracking procedure of Methods for detailed task description 1B: SERIA model. A reflexive early error is triggered if the early unit (dotted line) hits threshold first and is not inhibited by inhibitory unit. If inhibition is successful, action selection is a race between the late prosaccade unit (dashed line) and the antisaccade unit (solid line). The unit that hits threshold first determines the action (correct antisaccade or error) and the latency of the saccade. 1B adapted from (Aponte et al., 2017).

Given that desynchronization of beta band oscillations (13-30 Hz) is generally coined as a facilitator of movement initiation, its role during the preparation for saccade execution is also conceivable (Zhang et al., 2008). In fact, prefrontal beta power decreases in preparation for antisaccades contrasted with no-go trials in healthy individuals (Cordones et al., 2013). In an MEG study, while activity in the midfrontal areas was not analyzed due to low signal-to-noise ratio, increased beta band power was found over the lateral prefrontal cortex coupled with increased alpha band power over FEF during the preparation for antisaccades compared with prosaccades. Furthermore, higher pre-stimulus alpha power in FEF, which has been interpreted as a correlate of local inhibition, was associated with successful inhibition of a prepotent reflexive error (Hwang et al., 2014).

### Antisaccades in PD and the potential influence of DBS frequency

Patients suffering from PD tend to higher rates of erroneous prepotent saccades towards, instead of away from the stimulus, than healthy controls (Antoniades et al., 2015a; van Stockum et al., 2008; Waldthaler et al., 2019a), and the error rate has been found to correlate with executive dysfunction (Antoniades et al., 2015b). The influence of STN-DBS on response inhibition in the antisaccade task remains under debate. A recent meta-analysis of studies on the effects of STN-DBS on antisaccades in PD concluded that DBS reduces their latency significantly, while a moderate increasing effect on the antisaccade error rate did not reach significance, but was possibly underpowered with only five eligible studies (Waldthaler et al., 2021).

With 130 Hz as the default stimulation frequency, most patients are treated with DBS pulses between 60 and 200 Hz. A differential effect on several clinical hallmarks of PD has empirically evolved with higher frequencies enabling tremor control and lower frequency stimulation (60 to 90 Hz) possibly improving gait function and axial symptoms (Su et al., 2018). The limited evidence to date allows an application of low-frequency stimulation in individual cases (Conway et al., 2019). Nevertheless, further insight is warranted as higher DBS frequencies (>100 Hz) may have detrimental effects on cognition (Combs et al., 2015), while verbal fluency (Wojtecki et al., 2006) and cognitive interference (Varriale et al., 2018) may, on the contrary, even improve with low frequency pulses.

Interestingly, axial motor signs, and specifically freezing of gait, paralleled antisaccade performance in recent studies (Ewenczyk et al., 2017; Gallea et al., 2021; Nemanich & Earhart, 2016; Waldthaler et al., 2019b; Walton et al., 2015). Hallmark regions for gait impairment such as the pedunculopontine nucleus correlate in their functional connectivity with FEF, which in turn, correlates with antisaccade latency in an fMRI study (Ewenczyk et al., 2017; Gallea et al., 2021). Thus, a common underlying mechanism of freezing of gait and antisaccade control may be posit as an expression of a network-dependent degeneration (Ruppert et al., 2021). Given a possible modulation of gait with low-frequency stimulation (Moreau et al., 2008), the question seems pertinent whether 60 Hz-DBS may have an effect on antisaccades as well.

### Aim of this study and hypotheses

In this study, we combined eye-tracking, computational modelling of its behavioral outcomes and EEG recordings with the aim to explore effects of high (130 Hz)- vs. low-frequency (60 Hz) STN-DBS and no stimulation on response inhibition and its cortical correlates in the antisaccade task in PD-patients. The Stochastic Early Reaction, Inhibition, and late Action (SERIA) computational model allowed us to differentiate early and late antisaccade responses informing about underlying error mechanisms which might be attributed to either failures in response inhibition or in subsequent action selection (Figure 1B).

We hypothesized that:

1. Based on previous results, DBS at 130 Hz would result in more directive errors and decreases in latency of correct antisaccades as behavioral correlates of reduced response inhibition.
2. 130 Hz-DBS may reduce midfrontal theta power during the preparation for an antisaccade as an indicator for a stimulation-induced release from top-down cognitive control.
3. 60 Hz-DBS may have opposite effects to 130 Hz-DBS on response inhibition and midfrontal theta power, i.e., increasing midfrontal theta power and improving response inhibition.

Our hypotheses regarding 60 Hz-DBS should, however, be regarded as exploratory as they were based on a very limit number of studies reporting positive effects of 60 Hz-DBS on axial motor functions and cognition.

## METHODS

The study was approved by the Ethical Board of the University Hospital Marburg (reference number 119/19) and followed the Declaration of Helsinki. All participants gave written informed consent before participating. Patients were recruited from the Movement Disorders Outpatient Clinic of the Department of Neurology at the University Hospital Marburg.

### Participants

A sample size calculation can be found in the Supplementary Material 1. A total of 19 consecutive participants suffering from PD according to the clinical diagnostic criteria of the Movement Disorders Society (Postuma et al., 2015) and treated with chronic STN-DBS were recruited. All patients had undergone extensive monopolar review to find the optimal settings for DBS minimizing motor symptom and avoiding side effects. Pre-established exclusion criteria were 1) dementia according to the MDS task force criteria level 1 (Emre et al., 2007), 2) signs of clinically relevant depression (Beck Depression Inventory > 14 points), 3) history of other disorders of the CNS, 4) any concurrent conditions making eye-tracking or EEG recordings impossible (e.g., disorders of the eyes or visual system with reduced visual acuity, severe camptocormia, other orthopedic disorders impairing ability to sit for longer periods, etc.), and 5) medications possibly influencing eye movements or EEG recordings (in particular benzodiazepines).

Mean time between study inclusion and implantation of DBS leads was 10.2 ± 9.1 months with a minimum of three months to avoid any impact of lesion effects on the results. All participants were in off-medication state after overnight withdrawal of dopaminergic medication and at least 12 hours prior to the start of the assessments. Motor symptoms were rated on part III of the Movement Disorder Society – Unified Parkinson’s Disease Rating Scale (MDS-UPDRS) (Goetz et al., 2007). Levodopa equivalent daily doses were calculated according to (Tomlinson et al., 2010). Montreal Cognitive Assessment (MoCA) was used to evaluate general cognitive ability (Nasreddine et al., 2005).

The final data set included 14 participants. 8/19 participants had asked for pre-mature stopping of the study protocol (due to tiredness, unbearable motor symptoms or pain). Five of these had to be excluded from further data analysis because they did not complete at least one antisaccade block in all three conditions. While the remaining three participants ended the study prematurely, they had completed at least one block in each condition. Therefore, we decided to include their data in the further analyses. Please see Table 1 for a summary of demographic and clinical characteristics and Figure 2 for a visual summary of the study’s workflow.

**FIGURE 2:**
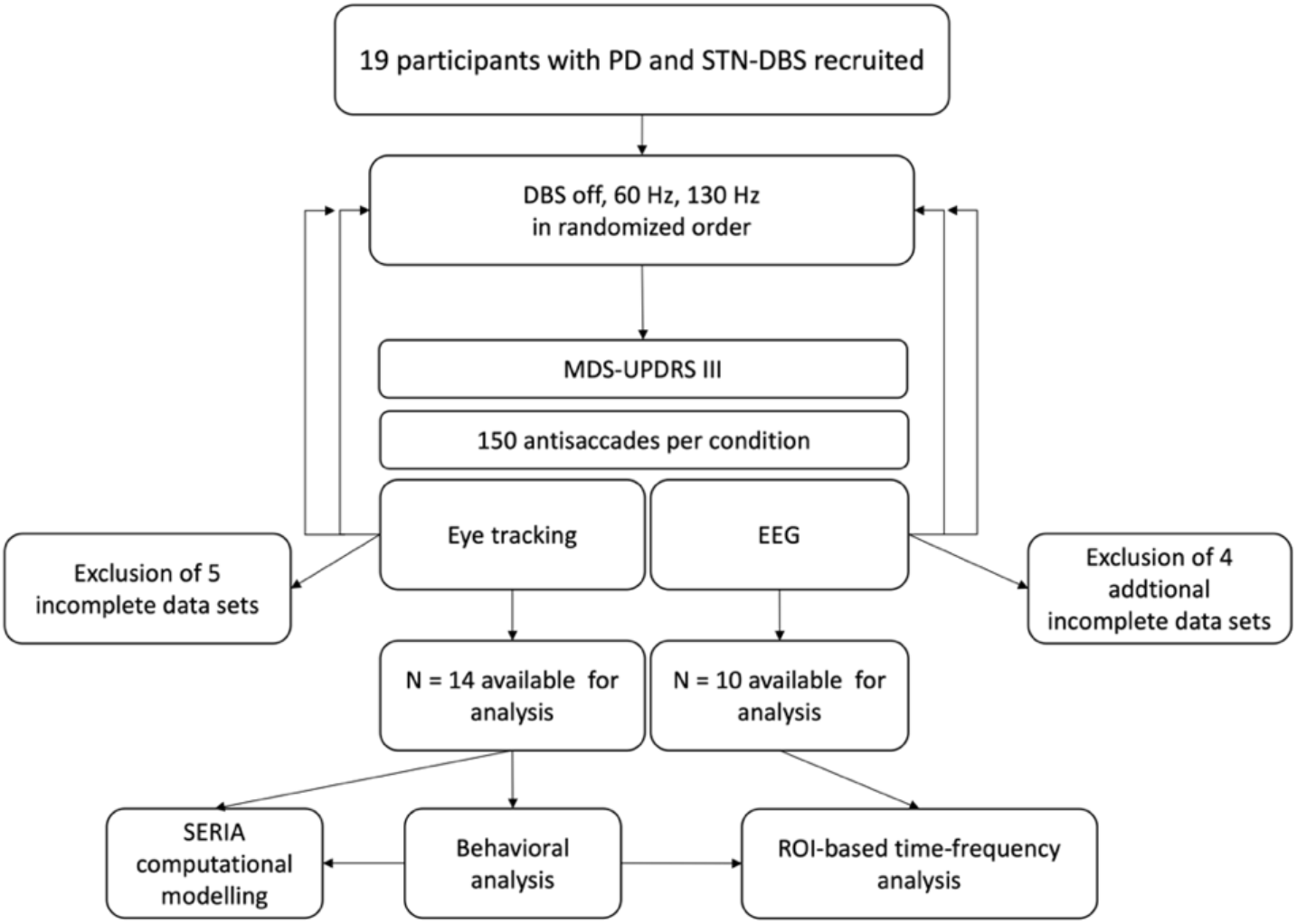
Study workflow. DBS = deep brain stimulation, MDS-UPDRS III = Movement Disorders Society Unified Parkinson’s Disease Rating Scale part III (motor), ROI = region of interest, SERIA = Stochastic Early Reaction, Inhibition, and Late Action model

**TABLE 1:**
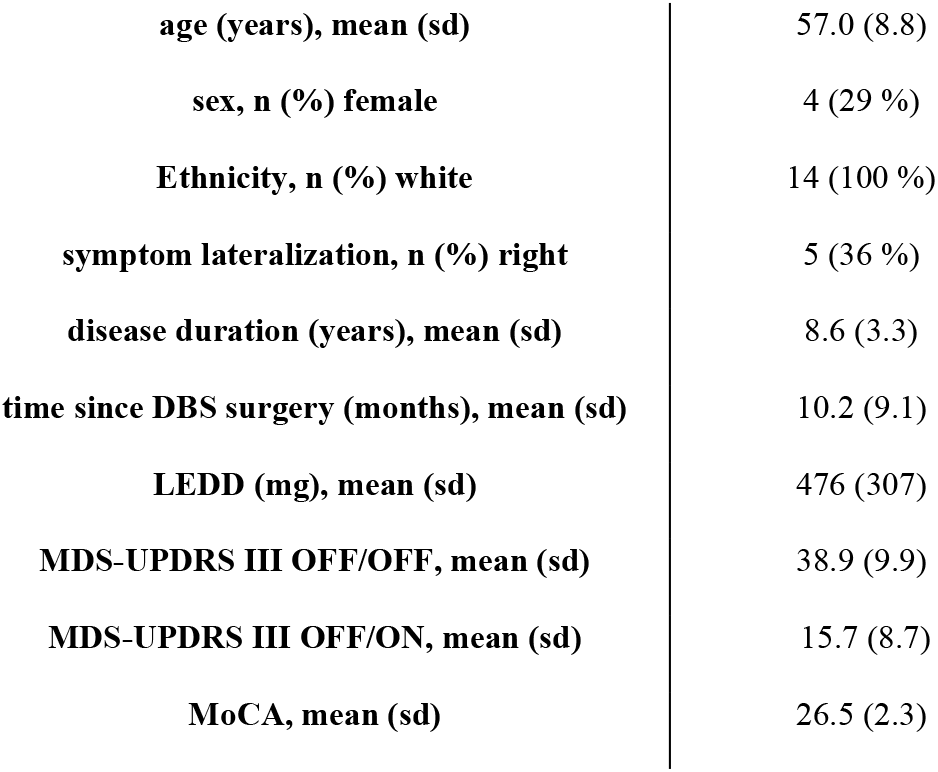
Demographics and clinical characteristics of the PD group

### DBS programming

Participants performed the task three times: i.) with DBS switched off, ii.) with DBS frequency set at 130 Hz and iii.) with DBS frequency set at 60 Hz. All other DBS parameters (contacts, amplitude, and impulse width) remained unaltered with respect to the chronic DBS program with optimal clinical response in each individual patient (cf. Supplementary Material 1). The participants were blinded for the active DBS program and there were wash-out periods between sessions of at least ten minutes. MDS-UPDRS III was assessed directly prior to the EEG recordings in each DBS condition.

There are recommendations to keep the total electrical energy delivered (TEED) constant between DBS programs which may be achieve by respective adjustments of stimulation amplitude (Moro et al., 2002). On the other hand, some authors discourage the use of TEED to censor or edit combinations of stimulation parameters (Marks, 2015). We decided against adjustments since the physiological role of TEED is subject of debate and increasing amplitudes in the 60 Hz condition (given that most participants were treated at higher frequencies) might have introduced additional bias or may have caused side effects (Koss et al., 2005).

### Eye-tracking procedure

Each participant completed recording sessions in all three DBS conditions on the same day. The order of conditions was randomized, and participants were blinded to avoid any biases due to expectation, learning effects or tiredness. The experiment took place in a sound-attenuated, darkened and electrically shielded room which the researcher monitoring the progress in the adjoining room. All participants were seated in an upright armchair with back support at distance of 70 cm from a computer monitor with a diagonal of 60 cm and with their head stabilized with chin and forehead rests.

An infrared video-based eye-tracker (EyeLink 1000 Plus, SR Research, Ontario, Canada) recorded positions of both eyes at a sampling rate of 500 Hz and an instrumental spatial resolution of 0.01° with simultaneous recording of EEG data. The eye-tracker was calibrated and validated with a 9-point grid before each experimental block. The validation was repeated until average errors for all points were <1° compared to the result of the calibration. Moreover, to ensure precision within blocks, a drift correction prior to each trial was performed.

The experiment was programmed in MATLAB 2020b (The Mathworks Inc., Massachusetts, USA) using the psychophysics toolbox (www.psychtoolbox.org) (Brainard, 1997). Three blocks of 50 horizontal antisaccades each were presented per condition (n = 150 per condition). Each trial started with a red central fixation cue (diameter 1° visual angle) that was presented for 1000 ms in the middle of a black screen. It was followed by the appearance of a white lateral target stimulus located either 10° left or right from the initial fixation cue (Figure 1A). The lateral stimulus was presented in equal numbers and random order to the left and right side of the screen. It vanished after 1000 ms and was followed by a white central dot for drift correction and a subsequent interstimulus interval (blank black screen) that allowed participants to blink. The next trial started with a new red central fixation cue.

The participants were instructed to look at the exact opposite direction of the lateral stimulus as fast and precisely as possible as soon as it was presented on the screen. Ten practice trials prior to the first antisaccade block of the experiment with verbal feedback ensured that participants understood the instructions. These practice trials were discarded. Between blocks, participants were given the opportunity to take breaks.

### Eye-tracking data processing and analysis

The researcher analyzing the eye-tracking and EEG data sets (JW) was not involved in data collection and was blinded to the participants’ identities. A parsing system incorporated in the EyeLink 1000 software intersected the raw eye position data into visual events, i.e., saccades, fixations, and blinks. This event data set was analyzed in the statistical computing program R (R Core Team, 2014) using the Eyelinker package. An acceleration larger than 8000°/s^2,^ peak velocity > 40°/s peak velocity, and a deflection > 0.1° were set as thresholds for saccade detection.

Saccade latency was defined as the time from stimulus onset to the start of the first saccade regardless of whether the saccade was elicited in the correct direction. A directive error was defined as a saccade towards lateral stimuli, i.e., a prosaccade. Saccades with latencies between 90 and 130 ms were defined as express saccades in the behavioral analysis and reported separately but were included in the computational model regardless of their direction.

Trials were removed from further analysis when i) the latency was in the anticipatory range (< 90 ms) or longer than two standard deviations from the individual mean latency of the participant, ii) the first saccade after stimulus onset had a starting position more than 3° lateral of central fixation dot, iii) a saccade with an amplitude smaller than 0.5 ° or larger than 15 ° was executed or iii) a blink occurred between stimulus presentation and the first saccade.

The following variables were defined as outcome measures: latency of correct antisaccades, latency of prosaccade errors, error rate (proportion of erroneous trials to all valid trials) and express saccade rate (proportion of express saccade trials to all valid trials). Processing of the eye-tracking data led to the rejection of a total of 16.8 % ± 11.5 % of trials (off: 19.7 % ± 14.3 %; 130 Hz: 14.4 % ± 11.3 %; 60 Hz: 16.2 % ± 8.6 %, χ2(2, 13) = 3.964, p = .138).

### Computational modeling of the eye-tracking data

Successful execution of antisaccades require response inhibition, i.e., withholding of an early, reflexive prosaccade, as well as the subsequent correct voluntary action selection. Thus, errors may occur when the early prosaccade is not stopped (*inhibition* error) or when the wrong action (i.e., a prosaccade) is selected later in the trial (*choice* error). However, these different types of errors cannot be directly measured based on the empirical eye-tracking data.

The Stochastic Early Reaction, Inhibition, and late Action (SERIA) model for antisaccades assumes that the latency and the response within trials stem from of a competition among four race-to-threshold processes or units: i) the early prosaccade unit, ii) the inhibitory unit (which inhibits an early prosaccade), iii) the antisaccade unit, and, iv) the late prosaccade unit (for details of the SERIA model cf (Aponte et al., 2017)). In brief, the parameters of SERIA capture the probability of an early inhibition failure as well as the probability of a late choice error and quantify the mean hit times of the early and the late units, i.e., the mean reaction times of early inhibition errors, correct antisaccades and late choice errors (Figure 1B).

The SERIA model was fitted using the open-source SEM toolbox (http://www.translationalneuromodeling.org/tapas/). The code was executed in MATLAB 2020b (The Mathworks, Inc., Massachusetts, USA) with GSL 2.7. A hierarchical method of fitting the model was applied to pool information across subjects. In this model, the prior distribution of the parameters of each subject was informed by the population distribution and thereby offers a form of regularization based on observations from the population. The parametric distributions for the increase rate (or reciprocal hit time) of each of the units was parametrized by a “mixed Gamma model”, such as the increase rate of the early and inhibitory unit was Gamma distributed, but the increase rate of the late units was inverse Gamma distributed. This decision was based on previous work by Aponte and colleagues demonstrating that an unconstrained mixed Gamma SERIA model was favored among a variety of different models (Aponte et al., 2018). The model was fitted using Markov chain Monte Carlo sampling via the Metropolis-Hastings algorithm. Evidence of marginal likelihood of the model was computed with thermodynamic integration with 16 chains and a 5th-order temperature schedule (Aponte et al., 2016). The algorithm was run for 15 × 10^4^ iterations with the first 6 × 10^4^ iterations being discarded as “burn-in” samples.

### EEG recording

EEG was recorded simultaneously during the eye-tracking sessions described above. We used an elastic cap with 128 electrodes mounted in a spherical array (Easy-Cap GmbH, Herrsching, Germany). To maintain electrode impedances below 10 kΩ, conduction gel was applied. The used caps were standardized and placed according to the 10/10 system. All data were recorded on a BrainAmp® standard amplifier (Brain Products GmbH, Gilching, Germany), low-pass filtered at 1 kHz and digitized at a sampling rate of 5 kHz. In addition to the scalp EEG electrodes, an electrocardiogram (ECG) electrode was placed for recording of cardiac activity.

### EEG preprocessing

EEG data processing and statistical analysis were run in MATLAB 2020b (The MathWorks Inc., Massachusetts, USA) and MNE Python (Gramfort et al., 2013) with Python version 3.7. First, data was resampled at 250 Hz, re-referenced to average and high-pass filtered to remove DC offset and drift (4^th^-order Butterworth filter, cut-off frequency 0.5 Hz). DBS artefacts were removed using the DBSFilt toolbox (https://github.com/guillaumelio/DBSFILT/blob/master/DBSFILT_GUI_DOC.pdf) which, briefly, filters the EEG signal and detects spikes based on the Hampel identifier for automated spike detection (Allen, 2009). This identifier treats artefacts as outliers in the frequency domain and replaces them with interpolated values, which was successfully used for DBS artefact removal before (Allen et al., 2010).

Additional EEG artefacts were detected and discarded as follows: First, bad channels were identified visually and corrected with the spherical spline method, which projects the sensor locations onto a unit sphere and interpolates the signal at the bad sensor locations based on the signals at surrounding artefact-free locations (Perrin et al., 1989). Consecutively, an independent component analysis (ICA) was used for blink as well as eye movement and heart artefact correction (3.2 ± 0.8 components removed) (Delorme et al., 2007).

Since this study focused on preparatory activity and eye movements were inherent in the response period of the trials, the epochs were limited to the time window before stimulus presentation, i.e., before the direction of the following saccade had been revealed to the participant. Thus, data were segmented into epochs time-locked to the onset of the cue stimulus containing the full 1000 ms period of fixation dot presentation (-1000 ms to 0 ms with respect to target onset). (Figure 1A). In this way, eye movement artefacts as well as any brain activity related to the sensorimotor transformation of the stimulus into a saccade (Moon et al., 2007) were excluded. Within the epochs’ time frame, the time window from 200 ms to 100 ms before presentation of the fixation dot was defined as baseline (-1200 ms to -1100 ms). The baseline was offset by 100 ms from fixation dot onset to minimize contamination of the baseline interval by fixation-associated activity. Only epochs in which a subsequent correct antisaccade was performed were further analyzed.

Four participants had to be excluded from the EEG analyses due to technical failure during the recordings resulting in a total of ten participants. Behavioral results are reported for the complete sample of 14 participants. The behavioral results in the subgroup that was included in the final EEG analysis did not differ from the entire sample.

In the remaining ten participants, trial rejections during eye-tracking and EEG preprocessing resulted in 49.7 ± 42.3 trials for DBS-off, 59.9 ± 42.1 trials for 130 Hz-DBS and 61.6 ± 29.8 trials for 60 Hz condition remaining for time-frequency analysis (F(2, 27) = 0.282, p = .8). For statistical testing, the number of trials was randomly equalized between all three conditions within each subject using the “equalize_epoch_counts” function implemented in MNE Python to maintain a constant signal-to-noise-ratio within subjects.

### Time-frequency analysis

Based on our *a priori* hypothesis, we restricted sensor-level EEG analyses to a selection of frontal EEG electrodes to avoid unnecessary multiple comparison. To focus on the hypothesized role of midfrontal theta oscillations during periods of enhanced cognitive control and during the preparatory period for an antisaccade in particular (Cordones et al., 2013; van Noordt et al., 2017; B. Zavala et al., 2016), time-frequency data from a midfrontal region of interest (ROI) encompassing the electrodes *F1, Fz* and *F2* were averaged. As right-lateralization of dynamics in DLPFC and iFG has been a recurrent finding in studies on response inhibition and antisaccade preparation (Hamm et al., 2012; Hwang et al., 2014; Swann et al., 2011), we also defined a right lateral prefrontal ROI including the electrodes AF8, F6, F8, and FC6. All further analyses were restricted to these two ROIs.

For each condition, time-frequency representations (TFR) of oscillatory power changes resulted from a Morlet wavelet decomposition with variable, frequency-dependent cycles (= frequency / 2) into frequency bins between 3 and 30 Hz. The length of the wavelets increased linearly from 1 cycle at 2 Hz to 15 cycles at 30 Hz to optimize the trade-off between temporal resolution at lower frequencies and stability at higher frequencies. The change in spectral power during the preparatory period (-1000 ms to 0 ms) is reported as the logratio from the baseline period (-1200 to -1100 ms from onset), calculated by dividing by the mean baseline power per frequency and converting to decibel (dB) by log-transformation (dB = 10 x log10(power/baseline), then averaged across trials for each condition.

To investigate effects of DBS conditions on cortical activity on a single-trial level, time-frequency transformations were adapted for single-trial analysis by calculating TFR for each trial separately using the same Morlet wavelet decomposition and baseline correction as outlined above. For mixed model logistic regressions that were used to predict trial outcome (correct antisaccade / error), TFR were additionally calculated for error trials, while only correct antisaccade trials were included in the single-trial mixed linear models for antisaccade latency. For the single-trial analysis, frequency ranges and time windows were determined post-hoc based on the group level results (see Results section).

### Statistical Analysis

The empirical outcomes (antisaccade latency, error latency, error rate and express rate) as well as the estimates of SERIA model (late saccade probability, late saccade reaction time, inhibitory fail probability, and inhibitory fail reaction time, antisaccade reaction time) were compared between conditions using one-way RM-ANOVA based on a general linear model (GLM) with subject as random effect in GraphPad Prism version 8.0.0 (GraphPad Software, San Diego, California, USA) Pairwise comparisons of the three conditions were performed using Tukey’s test with correction for multiple testing for normally distributed variables, respectively Friedman’s test followed by Dunn’s multiple comparison test for non-normally distributed variables. Normality was assessed with a Shapiro-Wilk test. Since we did expect differential effects of 130 Hz- and 60 Hz-DBS impulse frequency, we executed pairwise testing even if the RM-ANOVA resulted in an overall non-significant effect of condition. Statistical significance was asserted at α = 0.05.

Statistical inference of the EEG data was ascertained with cluster-based permutation tests implemented in MNE Python (Maris & Oostenveld, 2007). This approach corrects for multiple comparisons within time-frequency representations by identifying clusters of differences between conditions by summing adjacent significantly different time-frequency bins and comparing the cluster size to a distribution of largest cluster values obtained by randomly shuffling the conditional labels under the null-hypothesis. If the observed cluster statistics exceeded 95% of the permutation distribution (corresponding to critical α = 0.05) the null hypothesis was rejected. Cluster-based permutation one-way RM-ANOVA with 1000 permutations was used to compare the time-frequency representations between the three DBS conditions (off / 130 Hz / 60 Hz) followed by pairwise comparisons between the conditions using cluster-based permutation paired-t-tests with 511 permutations (exact full permutation test).

To assess whether preparatory beta or theta activity may predict the antisaccade outcome (correct antisaccade / error) on a single trial level, mixed model logistic regressions were run using the R package lme4 (Bates, 2005) with main effects of condition and theta, respectively beta activity (cf. Results section) as well as their interaction as fixed effects and participants as random effect. Further, mixed linear models were run to assess possible associations between preparatory EEG activity and antisaccade latency on a single trial level. Again, main effects of condition and theta, respectively beta activity as well as their interaction were entered as fixed effects into the model, while participants were treated as random effect. The resulting coefficients were deemed significant by the Satterthwaite approximation (Kuznetsova et al., 2016). P-values of the pairwise comparisons between the three conditions were Bonferroni-corrected to account for multiple comparison.

## RESULTS

### Behavioral results of the antisaccade task

No overall significant effect of condition (off / 60 Hz /130 Hz) on antisaccade latency (F(2,26) = 2.626, p = .1) was detected in RM-ANOVA. However, pairwise comparisons revealed that antisaccade latency was significantly decreased by -29.8 ms (CI = [6.3; 53.1]) with 130 Hz-DBS (q = 4.752, p = .01) compared with off-DBS state, while the mean reduction of 15.5 ms (CI = [-21.7; 52.7]) with 60 Hz-DBS was not statistically significant (q = 1.556, p = .5) (Figure 3E).

**FIGURE 3:**
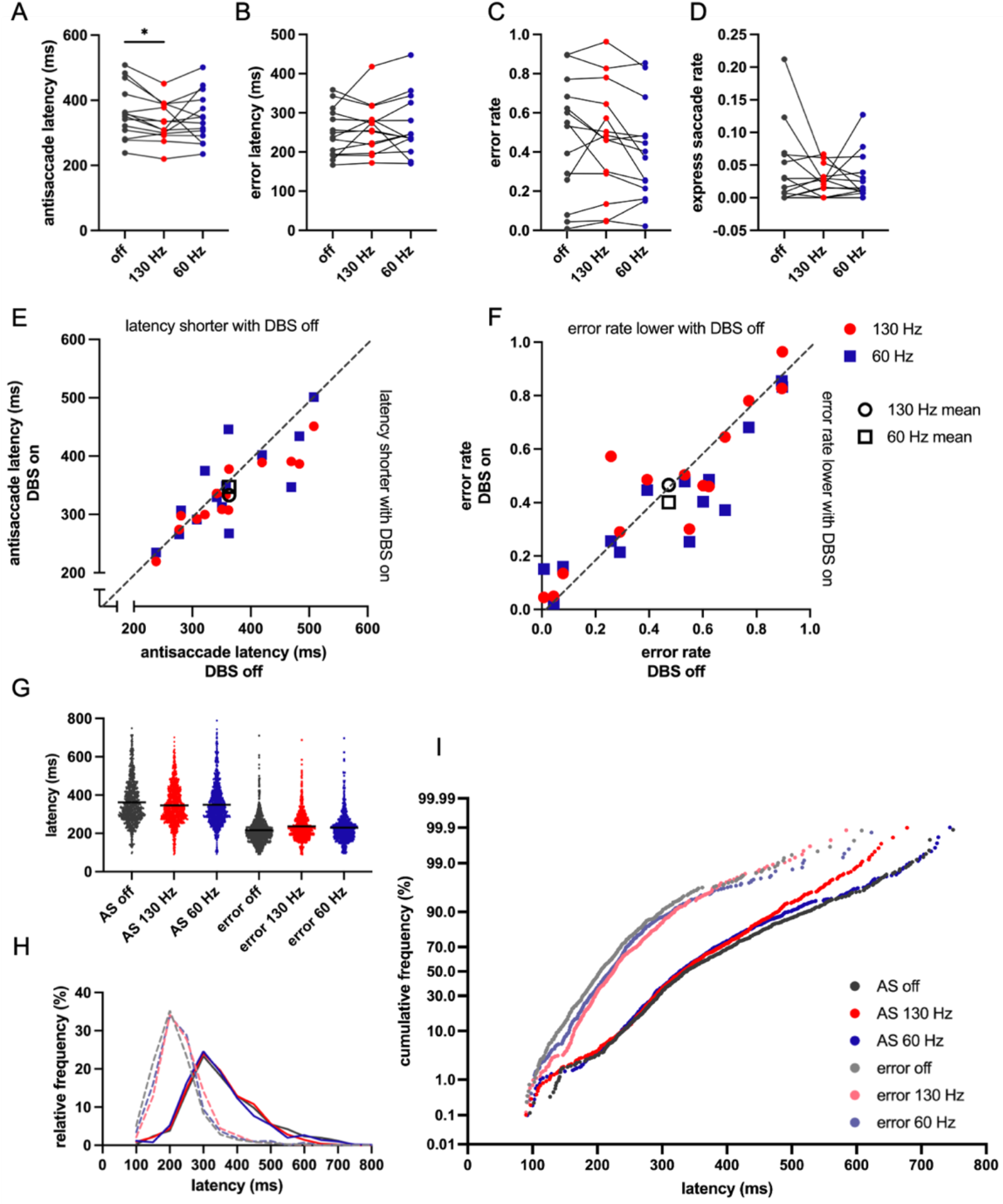
Behavioral results of the eye-tracking tasks. **A-D**: Antisaccade latency, error latency, error rate and express rate for each participant represented by an individual dot. Off-DBS state in gray, 130 Hz-DBS in red and 60 Hz-DBS in blue. Solid lines connect results from the same participant throughout conditions. * p<0.05. **E, F**: scatter plot comparing the mean of antisaccade latency (E), respectively error rate (F) in the DBS-off condition (x axis) and DBS-on conditions (y axis), individual values for each participant are displayed as red dots for 130 Hz-DBS and as blue squares for 60 Hz-DBS, the black dot and square represent the group means. The unity line is shown as a dark gray dashed line. **G-I**: Latency frequency distributions. **G**: Dot plots showing latencies of all antisaccade (AS), and error trials pooled for all participants. Black horizontal lines represent the mean. **H**: Relative frequency distributions of latencies in bins of 50 ms. Solid lines represent correct antisaccade trials, dashed lines represent errors. **I**: Cumulative frequency distributions of latencies with probit scaled y axis. Correct antisaccades (AS) displayed in intense colors and errors in faint colors. Individual data for each participant that was used to create these plots are available in Figure 3 – source data 1.

No significant between-condition differences were detected in the RM-ANOVA nor in pairwise comparisons for error latency (F(2,26) = 1.118, p = .3), error rate (F(2,26) = 2.790, p = .08) and express rate (χ2(2, n = 14) = 0.154, p = .9) (Figure 3, Table 2).

**TABLE 2:** Behavioral results of the antisaccade task and predicted outcomes of the SERIA model. To emphasize the difference between the variables, empirical reaction times are referred to as latencies and model-based reaction times are referred to as hit times. Significant differences between the conditions are marked in bold / red. ^1^ – variable did not pass Shapiro Wilk test, non-parametric testing (Friedman test followed by Dunn’s test) was applied, and rank sum differences are reported instead of mean differences. CI – confidence interval, diff – difference, padj – adjusted p value after correction for multiple testing, sd – standard deviation, se – standard error

|  | off-DBS |  | 130 Hz-DBS |  | 60 Hz-DBS |  | off vs. 130 Hz |  |  |  | off vs. 60 Hz |  |  |  | 130 Hz vs. 60 Hz |  |  |
| --- | --- | --- | --- | --- | --- | --- | --- | --- | --- | --- | --- | --- | --- | --- | --- | --- | --- |
|  | mean | sd | mean | sd | mean | sd | mean<br>diff. | se of<br>diff. | 95% - CI | p<sub>adj</sub> | mean<br>diff. | se of<br>diff. | 95% - CI | p<sub>adj</sub> | mean<br>diff. | se of<br>diff. | 95% - CI |
| **empirical outcomes** |  |  |  |  |  |  |  |  |  |  |  |  |  |  |  |  |  |
| antisaccade |  |  |  |  |  |  |  |  |  |  |  |  |  |  |  |  |  |
| latency | **362.9** | **80.9** | **333.1** | **60.1** | **347.4** | **76.7** | **29.8** | **8.9** | **6.3, 53.1** | **0.013** | 15.5 | 14.1 | -21.7, 52.7 | 0.531 | -14.2 | 15.1 | -54.2, 25.7 |
| error latency | 250.7 | 61.3 | 262.6 | 63.6 | 266.6 | 77.2 | -11.8 | 11.5 | -42.2, 18.5 | 0.572 | -15.8 | 10.5 | -43.7, 12.0 | 0.321 | -4.0 | 11.0 | -33.0, 25.0 |
| error rate | 0.473 | 0.301 | 0.466 | 0.283 | 0.401 | 0.252 | 0.007 | 0.036 | -0.087, 0.101 | 0.977 | 0.073 | 0.035 | -0.020, 0.165 | 0.134 | 0.065 | 0.031 | -0.016, 0.146 |
| express rate<sup>1</sup> | 0.044 | 0.060 | 0.025 | 0.023 | 0.030 | 0.036 | 1<sup>1</sup> |  |  | >0.999 | -1<sup>1</sup> |  |  | >0.999 | -2<sup>1</sup> |  |  |
| **SERIA model** |  |  |  |  |  |  |  |  |  |  |  |  |  |  |  |  |  |
| probability of |  |  |  |  |  |  |  |  |  |  |  |  |  |  | - |  |  |
| early error | **0.391** | **0.250** | **0.268** | **0.202** | **0.278** | **0.245** | **0.123** | **0.039** | **0.020, 0.226** | **0.020** | **0.114** | **0.040** | **0.008, 0.220** | **0.035** | 0.009 | 0.046 | -0.130, 0.112 |
| probability of late |  |  |  |  |  |  |  |  |  |  |  |  |  |  |  |  |  |
| error<sup>1</sup> | **0.179** | **0.232** | **0.272** | **0.268** | **0.168** | **0.204** | -10.0<sup>1</sup> |  |  | 0.176 | 4.0<sup>1</sup> |  |  | >0.999 | **14.0<sup>1</sup>** |  |  |
| antisaccade unit |  |  |  |  |  |  |  |  |  |  |  |  |  |  |  |  |  |
| hit time | 372.2 | 89.2 | 335.1 | 61.7 | 347.1 | 85.0 | 37.1 | 15.3 | -3.4, 77.5 | 0.074 | 25.1 | 17.1 | -20.1, 70.3 | 0.337 | -12.0 | 27.1 | -57.9, 34.0 |
| inhibitory unit hit |  |  |  |  |  |  |  |  |  |  |  |  |  |  |  |  |  |
| time | 211.9 | 54.5 | 236.4 | 60.3 | 221.9 | 65.8 | -24.5 | 20.2 | -77.9, 29.0 | 0.469 | -9.9 | 19.5 | -61.5, 41.6 | 0.869 | 14.5 | 18.5 | -34.3, 63.4 |
| late prosaccade |  |  |  |  |  |  |  |  |  |  |  |  |  |  |  |  |  |
| unit hit time | **327.7** | **87.4** | **282.3** | **50.1** | **311.8** | **74.6** | **45.3** | **14.8** | **6.2, 84.4** | **0.023** | 15.9 | 17.2 | -29.5, 61.3 | 0.635 | -29.4 | 17.3 | -75.2, 16.3 |

To further visualize the distribution of latencies across trials, the relative and cumulative latency distributions of all trials are shown color-coded for the three conditions (Figure 3G-I). Although not statistically significant in the RM-ANOVA, a prominent feature in the off-DBS condition compared to 130 Hz and 60 Hz-DBS was a high proportion of early error saccades, including saccades within the express (<130 ms) and very fast reflexive (< 150 ms) range .

### SERIA model of antisaccades

As expected, the behavioral analysis revealed a decreasing effect of 130 Hz-DBS on antisaccade latency but did not show the hypothesized increasing effect on antisaccade errors. In contrast, the antisaccade error rate was reduced by switching 60 Hz-DBS on in 10 of 14 participants (Figure 3F) with a mean reduction of 7.3 % compared to off-DBS state, although without significant group effect. To relate these behavioral findings to differences between the conditions, the SERIA model was applied to the empirical data. The main aim was to determine whether different DBS frequencies differentially affected 1) the hit time of the inhibitory (response inhibition) and late units (action selection), 2) the probability of inhibition failures (response inhibition), and 3) the probability of late choice errors (action selection, see Methods section).

In off-DBS state, the proportion of early inhibition failures was estimated to be 39.1 ± 25.0 % of all trials, respectively 69.5 ± 21.9 % of all errors. Accordingly, 17.9 ± 23.2 % of all trials, respectively 30.5 ± 21.9 % of errors were considered late prosaccade errors.

The effect of condition on the probability of errors due to early inhibition failure was significant in RM-ANOVA (F(2,26) = 5.364, p = 0.01). Pairwise comparisons revealed a significant reduction of the probability of early inhibition errors by 12.3 % (95%-CI = [2.0, 22.6] with 130 Hz-DBS (q = 4.451, p = .02) and by 11.4 % (95%-CI = [0.8, 22.0] with 60 Hz-DBS (q = 4.005, p = .04) compared with off-DBS state (Figure 4A).

**FIGURE 4:**
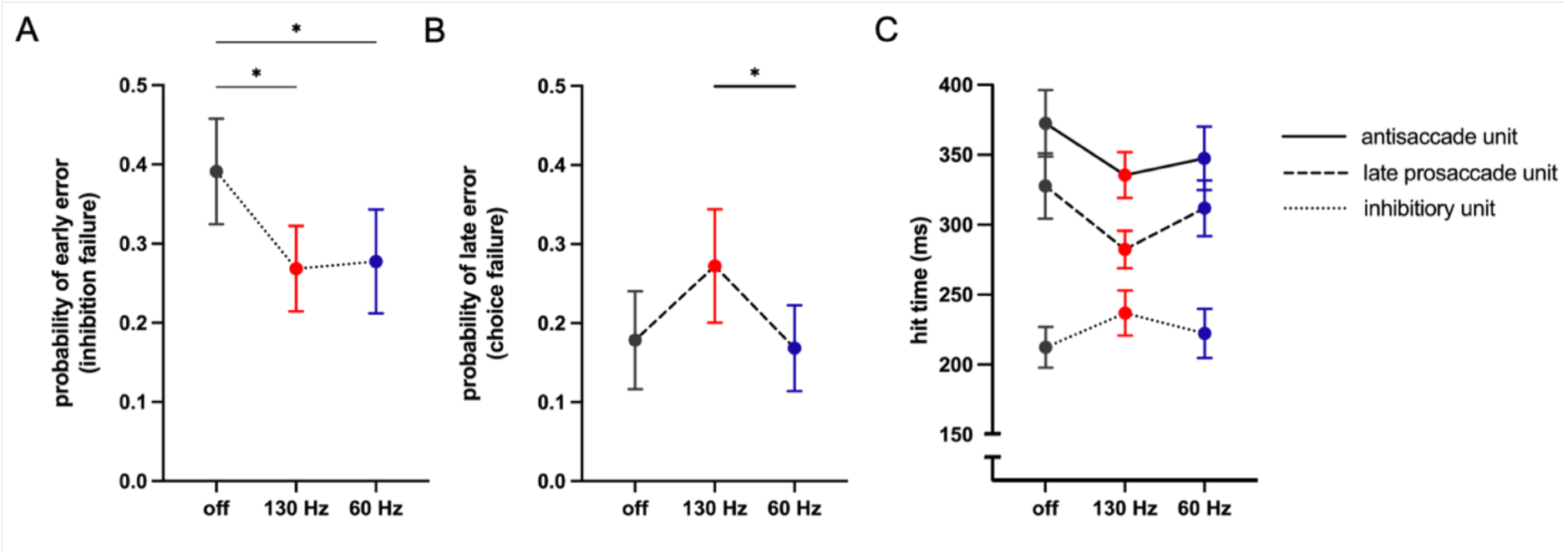
Results of the SERIA modeling of antisaccade performance. Probabilities for an early inhibitory (3A) and late choice error (3B) and hit times of the early prosaccade unit, late prosaccade unit and antisaccade unit (3C) as predicted by the SERIA model. Off-DBS state in gray, 130 Hz-DBS in red and 60 Hz-DBS in blue with dots representing the mean and vertical lines representing the standard error. The lines connecting the dots indicate the unit with solid lines representing the antisaccade unit, dashed lines representing the late prosaccade unit and dotted lines representing the early prosaccade unit (i.e., failure of the inhibitory unit). * p < .05 (not shown in C, please see main text). Individual data for each participant that was used to create these plots are available in Figure 4 – source data 1. SERIA model results per participant and correlations between behavioral outcomes of the antisaccade task and SERIA predictions are available in Figure 4 - Supplement 1.

A Friedman test indicated a significant effect of the DBS condition on the probability of late prosaccades (χ2(2, n = 14) = 7.429, p = .02) with pairwise comparisons resulting in a significant increase by 10.4 % with 130 Hz-DBS compared with 60 Hz-DBS (Z = 2.646, p = .02, see also Figure 4B)

Regarding the expected hit times, RM-ANOVA showed that the late prosaccade unit was significantly affected by condition (F(2,26) = 3.893, p = .04). Pairwise comparisons revealed that this effect was driven by a significant mean reduction of hit time of late prosaccades with 130 Hz-DBS by 45.3 ms (CI = [6.2, 84.4], q = 4.330, p = .02) compared with off-DBS state (Figure 4C).

There was no significant overall effect of condition on the hit time of the antisaccade unit (F(2,26) = 2.588, p = .1). In pairwise comparisons, a mean reduction by 37.1 ms (CI = [-3.4, 77.5], q = 3.423, p = .07) with 130 Hz-DBS compared with off-DBS state did not reach statistical significance, however with most of the confidence interval indicating in the direction of a decrease in hit time with 130 Hz-DBS.

There was no significant effect of condition on the hit times of the early inhibitory unit in the RM-ANOVA (F(2,26) = 0.802, p = .5) nor in pairwise comparisons. For complete results please also see Table 2.

To evaluate the validity of the SERIA model, we assessed the correlations between empirical and predicted error rates, antisaccade latencies and error latencies which resulted in high correlations coefficients ranging from 0.94 to 0.99 for all DBS conditions (Figure 4 - Supplement 1.).

Taken together, the results of the computational model of antisaccade performance suggest that the DBS condition had a differential effect based on its frequency:

1. 130 Hz and 60 Hz-DBS decreased the probability of an early inhibition failure compared with off-DBS state, indicating improved response inhibition early in the trial.
2. 130 Hz-DBS induced a reduction of the hit time of the late prosaccade unit and a trend towards a reduction of the hit time of the antisaccade unit compared with off-DBS state accompanied by an increased probability of a late prosaccade error compared with 60 Hz-DBS, indicating faster but more error prone action selection later in the trial.

### Preparatory midfrontal EEG dynamics

The results of the one-way RM-ANOVA with permutation clustering comparing the TFR in the midfrontal ROI between the three DBS conditions are presented in Figure 5. RM-ANOVA showed a significant cluster indicating a main effect on beta power (18 – 22 Hz) in the midfrontal ROI during the early preparatory period (p = 0.04) (Figure 5B). Pairwise comparisons revealed that this effect was driven by a larger decrease in beta power (18-26 Hz) from baseline between -1000 ms and approximately -500 ms in 130 Hz-DBS (p = 0.01) and between -1000 ms and approximately -300 ms in 60 Hz-DBS (p = 0.04) compared with off-DBS state (Figure 5C-E).

**FIGURE 5:**
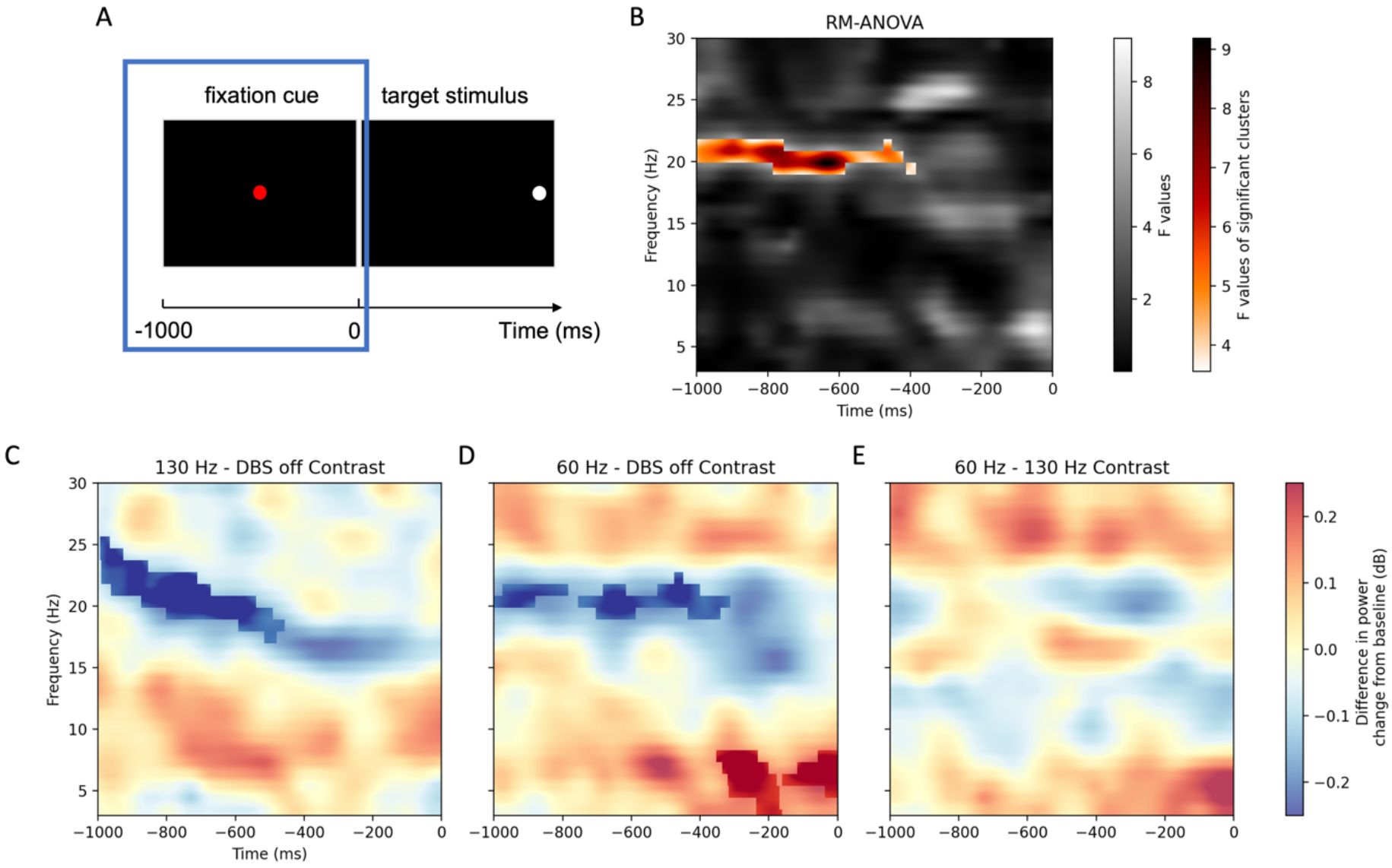
A: The time window of the TFR analysis corresponds to the presentation of the fixation cue during the antisaccade trial. B: Results of the RM-ANOVA comparing the averaged time-frequency representations in the midfrontal region of interest (Fz, F1, F2) between the three DBS conditions. Highlighted is the cluster of significant differences in power change that led to the rejection of the null hypothesis. Non-significant F values in sequential gray, F values corresponding to the cluster of significant group difference in color. C-E: Time-frequency representations of the contrast between conditions. Bold colors highlight the significant clusters in the pairwise comparisons. Null-results of the same analysis of preparatory EEG dynamics for the lateral prefrontal ROI is available in Figure 5 – Figure Supplement 1.

A second significant cluster indicating a theta effect (4 – 8 Hz) during the second half of the preparatory period from approximately -400 ms to stimulus onset at 0 ms was observed in pairwise comparisons between the 60 Hz-DBS condition and DBS-off state (p=0.04) (Figure 5D).

For the lateral prefrontal ROI, RM-ANOVA with permutation clustering resulted in no significant differences of TFR between DBS conditions (Figure 5 – Figure Supplement 1).

### Condition-dependent single-trial predictive value of midfrontal theta power

Based on the results above, we restricted the single-trial analysis to the frequency ranges (beta: 18-26 Hz, theta: 4-8 Hz) and time windows (beta: -1000 ms to -300 ms, theta: -400 ms to 0 ms) for which significant effects of DBS condition were identified in the group level analysis.

In the linear mixed model evaluating the relationship between midfrontal theta power, DBS condition and antisaccade latency with participants as random effect, we observed a main effect of condition on antisaccade latency as expected from behavioral findings (χ^2^(2) = 32.397, p < 0.001) with significant differences between 130 Hz-DBS and off-DBS state (β = 0.238, 95%- CI = [0.15, 0.32], t(1654) = 5.565, p_adj_ < 0.001) and between 130 Hz-DBS and 60 Hz-DBS (β = 0.164, 95%-CI = [0.09, 0.24], t(1654)= 4.112, p_adj_ < 0.001). There was no main effect of theta power on antisaccade latency (χ^2^(1) = 0.942, p = 0.3). However, the interaction effect between condition and theta power was found to be significant (χ^2^(2) = 7.327, p = 0.03), with 130-Hz DBS differing from the off-DBS state (β = 0.112, 95%-CI = [0.03, 0.19], t(1654) = 2.703, p_adj_ = 0.01), indicating that the effect of theta activity on antisaccade latency varies between these two conditions. From Figure 6, it is evident that as theta increased, antisaccade latency increased in off-DBS state, while it decreased with 130 Hz-DBS. Thus, 130 Hz-DBS reversed the effect of midfrontal theta activity on antisaccade latency.

**FIGURE 6:**
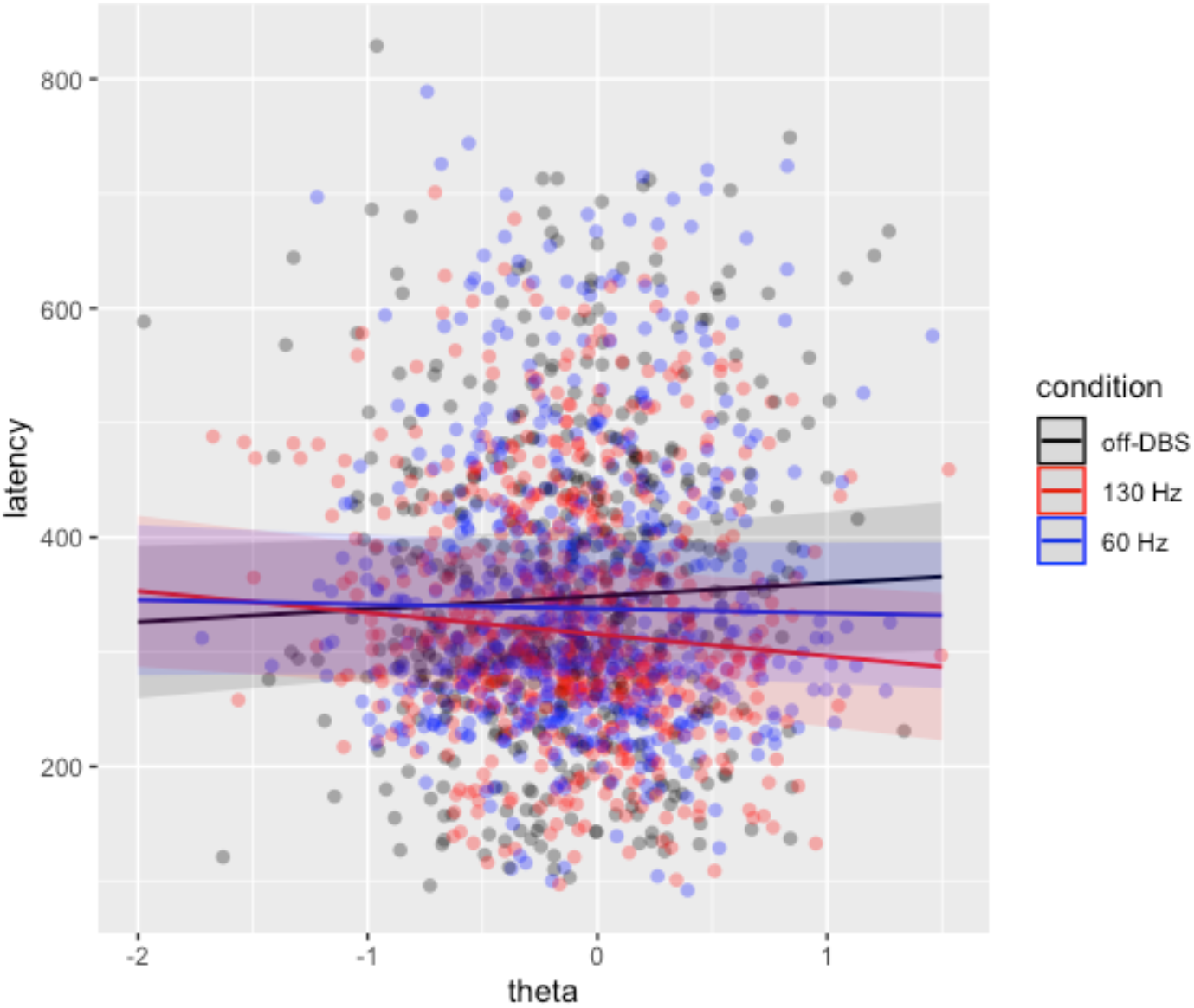
Single-trial mixed linear regression model of the relationship between antisaccade latency (in ms), midfrontal-theta activity (in dB) and DBS condition (off-DBS in black, 130 Hz DBS in red and 60 Hz-DBS in blue) as fixed effects and participants as random effect. Dots represent single trials. Shaded areas represent 95%-confidence intervals. The theta x condition interaction differed significantly between off-DBS and 130 Hz DBS. Raw trial data that was used to create this graph is available in Figure 6 – source data 1.

No significant effects of cortical power nor of power x condition interactions were identified in the linear mixed model including beta power as well as in the mixed logistic regression models of error probability. For complete results, please see Supplementary Material 2.

## DISCUSSION

### Summary of findings

In this study, we aimed at comparing the effects of high- and low-frequency STN-DBS on response inhibition in PD-patients using the antisaccade task in combination with EEG recordings. Despite no statistically significant effect of DBS on the total antisaccade error rate, computational modelling revealed that the probability of an early error due to failed inhibition of a prepotent reflexive prosaccade decreased with both 130 Hz and 60 Hz-DBS compared with the off-DBS state. Against our a priori hypothesis, these findings suggest that STN-DBS may *improve* response inhibition in PD.

Given the short latency of early errors allowing very limited processing time after stimulus presentation, these early inhibition failures most likely result from a lack of proactive response inhibition prior to stimulus onset during the mental preparation period for the task (Aponte et al., 2017). In the EEG analysis of this preparatory period before stimulus presentation, high and low frequency STN-DBS both induced stronger decreases of midfrontal beta power compared with off-DBS state. As attenuation of beta oscillations is theorized to support movement initiation (Engel & Fries, 2010; Y. Zhang et al., 2008), enhanced beta desynchronization with 60 Hz and 130 Hz-DBS may reflect a proactive modulating effects on oculomotor brain areas.

Yet, most strikingly, midfrontal theta band power increased during the preparatory period exclusively in 60 Hz-DBS compared with off-DBS state which may be considered an EEG correlate of enhanced cognitive control with low-frequency pulses. This may be supported by the fact that in trial-by-trial analyses higher preparatory midfrontal theta power predicted longer latencies of the upcoming saccade in off-DBS state which is consistent with studies reporting an association of higher midfrontal theta power with increased response times in various cognitive tasks (Cohen & Cavanagh, 2011; Cooper et al., 2019; van Driel et al., 2015). In line with previous evidence (Bakhtiari et al., 2020; Rivaud-Péchoux et al., 2000; Yugeta et al., 2010), antisaccade latency decreased when 130 Hz-DBS was switched on, while there was no behavioral effect of 60 Hz-DBS on antisaccade latency. Computational modelling revealed that the acceleration in the 130 Hz-DBS state was most likely due to faster, but more error-prone action selection processes later in the trial as indicated by decreased hit times of the late prosaccade unit and the antisaccade unit that were accompanied by an increased probability of late prosaccade errors. Interestingly, 130 Hz-DBS also reversed the relationship between midfrontal theta activity and antisaccade latency, indicating that high frequency STN-DBS may disrupt aspects of theta-based preparatory cognitive control.

### Attenuation of preparatory beta power as a proactive mechanism

Since beta desynchronization is generally coined as facilitator of movement initiation and of changing the ongoing motor set, the attenuated pre-stimulus beta activity under DBS pulses at 130 and 60 Hz alike suggests higher levels of early proactive activation of the oculomotor network regardless of stimulation frequency. In line with this interpretation, healthy individuals also show prefrontal pre-stimulus beta desynchronization in antisaccades when contrasted with no-go trials (Cordones et al., 2013). As such, our data is consistent with current theories postulating an anticipatory, proactive role of beta power modulation in the preparation for motor and cognitive responses (Jenkinson & Brown, 2011; Oswal et al., 2012). In PD, a lack of preparatory beta desynchronization has been interpreted as a general shift from proactive to more reactive motor control (Praamstra & Pope, 2007; Te Woerd et al., 2015). Consistent with our findings, STN-DBS may attenuate aberrant cortical beta activity (Abbasi et al., 2018; Devos et al., 2004).

At the same time, stability of beta oscillations also facilitates motor inhibition, so that its attenuation may also result in a higher probability of errors. In this regard, Hamm and colleagues found that beta power in ACC was lower for errors than for correct antisaccade trials in healthy individuals (Hamm et al., 2012). The authors argued that tonic beta activity may be crucial for correct antisaccade execution as it prevents errors by maintaining the ongoing oculomotor set, i.e., fixation instead of an early reflexive saccade. Notably, enhanced beta desynchronization with STN-DBS in our study was found in trials with a subsequent successful antisaccade. Thus, a certain level of beta suppression might be necessary to permit the dynamic reconfiguration of neural networks into a state of readiness for executive processing (Oswal et al., 2012). Furthermore, there was no significant detrimental behavioral effect of STN-DBS on antisaccade error rates, nor did the computational model suggest increased inhibition failures (i.e., early errors), but even the opposite. Therefore, we hypothesize that STN-DBS normalized the amount of preparatory beta desynchronization, which had been diminished in the off-medication and off-DBS state in the PD cohort (Singh, 2018), allowing sufficient proactive preparation of the oculomotor network without causing impulsive early responses.

Moreover, successful response inhibition may only be associated with a subsequent increase in prefrontal beta power after the target has been presented, but not during the cue period (Liebrand et al., 2017). Thus, the relationship between prefrontal beta activity and antisaccade outcome may differ after the target direction has been revealed. Since we did not analyze the changes in cortical oscillations after target presentation, any further considerations on potential changes of beta power later in the trial are beyond the scope of this study.

### Midfrontal theta power and response inhibition

Midfrontal theta activity reflects cortical correlates of cognitive control which may be exerted via synchronized activity in a prefrontal-subthalamic network (B. Zavala et al., 2016). It has been proposed that medial frontal cortical areas, e.g. the ACC, activate the STN to inhibit impulsive actions via theta oscillations as soon as conflicts arise or the need for cognitive control is detected (B. A. Zavala et al., 2014). In healthy individuals, error trials, as compared to correct antisaccades, were associated with a lack of increase in midfrontal theta during the preparatory period (van Noordt et al., 2017).

Further, ACC has been shown to have top-down control over the frontoparietal oculomotor network during the preparatory period for antisaccades, supported by a strong theta and beta synchronization from ACC to FEF (Babapoor-Farrokhran et al., 2017). Together, these preparatory oscillatory changes may subsequently prevent an early reflexive prosaccade (that is an error) when the stimulus is presented, thereby allowing additional time needed to activate the correct oculomotor set for a voluntary saccade later in the trial. As 60 Hz-DBS increased preparatory midfrontal theta activity and reduced the probability of an early reflexive error, our results support that 60 Hz STN-DBS may improve response inhibition by enhancing proactive cognitive control in PD via midfrontal theta oscillations.

### The influence of STN-DBS on midfrontal theta power and antisaccade latency

In our single-trial EEG analysis, higher theta activity during the late preparatory phase precited longer antisaccade latency in the off-DBS state. Consistent with this finding, theta activity has been associated with a slowing of the upcoming response in a variety of cognitively demanding tasks (Cohen & Cavanagh, 2011; Cooper et al., 2019; van Driel et al., 2015). Given this relationship in healthy controls and individuals with PD in off-DBS state, one may also expect an increase in antisaccade latency with the increase in midfrontal theta power with 60 Hz-DBS which was, however, not supported by our data.

Conversely, the trial-by-trial correlation analysis revealed an interesting inversion of the relationship between midfrontal theta power and antisaccade latency with 130 Hz-DBS. While surprising at first glance, this finding is in line with an influential study by Cavanagh and colleagues who found the same inversion of the relationship between midfrontal theta and response times in a decision-making task when STN-DBS was switched on. (Cavanagh et al., 2011). The authors concluded that STN-DBS disrupted the functionality of the medial prefrontal – STN network, which, if intact, would raise the decision threshold. Thereby, the impact of parallel cortico-striatal mechanisms, e.g., via pre-SMA and striatum, might increase which would facilitate high-value actions and reduce decision thresholds (Forstmann et al., 2008).

In the model of striatal action selection, the STN is pivotal for inhibiting prepotent actions under conflict (Zaghloul et al., 2012), i.e., when more than one potential response set are triggered simultaneously and compete to be selected as a response to the same external stimulus (here: a correct antisaccade versus a late visually guided prosaccade error) (Herz et al., 2018). 130 Hz-DBS may interfere with the delaying impact of STN on this “race” between the competing inputs (Jahanshahi et al., 2015) by disruption of theta-mediated pathways between midfrontal regions and STN. That 130 Hz-DBS may indeed alter the speed-accuracy trade-off with faster, but more error-prone action selection was supported by the computational model of the behavioral data showing an acceleration of late responses with 130 Hz-DBS. The exact mechanism that underlies late responses in the SERIA model is, however, an area of speculation. It seems plausible that processes after revelation of the stimulus direction exert a main role, e.g., the selection of an appropriate action and the subsequent sensorimotor transformation into a saccadic eye movement (Aponte et al., 2017).

Both the reversing effect on the relationship between midfrontal theta and antisaccade latency as well as the effects on late responses in the computational model were specific for high-frequency DBS. In particular, the chance for a late choice error was significantly increased with 130 Hz-DBS compared with 60 Hz-DBS, indicating that 130 Hz-DBS, but not 60 Hz may facilitate impulsive choices later in the trial.

### Limitations and future directions

A major limitation of our study is its comparatively small sample size. However, recruitment of eligible participants with PD and STN-DBS without any exclusion criteria is inherently very limited. Additionally, the study protocol was challenging to complete for this population as supported by the high proportion of pre-mature withdrawals of 44 % despite careful screening of potential participants.

Since we included no healthy control group, we cannot state whether response inhibition was overall impaired in the PD group. However, the mean antisaccade error rate of 47.3 % in off DBS-state is within the range of comparable studies in PD and considerable higher than in healthy age-matched controls (Waldthaler et al., 2021).

Participants completed the study in off-medication state. While this is a clear advantage of the study since it excludes effects of dopaminergic medication on the results to a large extent (long lasting effects > 12 hours cannot be entirely excluded), dopamine replacement therapy and DBS may interact in their effects on impulsivity in PD in real life scenarios. For instance, Bakhtari and colleagues showed that dopamine replacement therapy partly restored the detrimental effect of STN-DBS on antisaccade error rates (Bakhtiari et al., 2020).

Participants were stimulated with their individual optimal DBS program and amplitude, impulse width, and DBS contacts were not changed for the study. Thus, DBS settings were not standardized between participants. On the other hand, standardization of DBS settings would have carried a high risk for side effects since therapeutic and side effects of DBS vary widely between patients and optimal settings are the result of highly individualized programming procedures. Further, additional factors such as individual deviations from optimal lead placement could have not been standardized anyway. By keeping the individualized optimal DBS settings instead (other than frequency), we aimed to avoid side effects and to resemble the DBS effect achieved in daily life with chronic stimulation.

TEED was not kept constant between the DBS settings used in the study. TEED is expected to be lower with 60 Hz than with 130 Hz stimulation when stimulation amplitude is kept constant. As a recent study showed that changing DBS amplitude influences antisaccade performance (Munoz et al., 2021), we cannot exclude that any performance differences may be related to TEED differences (see also Methods section for further elaboration on this issue).

## CONCLUSION

In summary, a combined approach including eye-tracking, computational modelling and EEG allowed us to differentiate the effects of two commonly used STN-DBS frequencies on early and late responses in the antisaccade task and to explore their cortical correlates.

While 130 Hz-DBS may improve response inhibition and, thereby, reduce early impulsive actions, it seems to induce an altered speed-accuracy trade-off resulting in a higher likelihood for later impulsive choices. Since 130 Hz DBS reversed the relationship between midfrontal theta activity and antisaccade latency, it may disrupt theta-mediated medial prefrontal-STN interactions and thereby interfere with action selection processes.

60 Hz-DBS may provide the beneficial effect on response inhibition accompanied with an increase in midfrontal theta power without causing the detrimental effect on action selection and later responses seen with 130 Hz DBS. Here, our results warrant future studies on the cognitive effects of low-frequency STN-DBS in PD.

Furthermore, inconclusive behavioral results of previous and upcoming studies on the effects of STN-DBS on cognitive control should be interpreted in the light of potentially opposing effects of different DBS frequencies on various aspects of impulsivity in PD.

## Source Data Files

Figure 3 – Source Data 1: Averaged data per participant used to create this figure

Figure 4 – Source Data 1: Results of the SERIA model per participant used to create this figure

Figure 6 – Source Data 1: Trial-wise eye-tracking data and theta power over the midfrontal ROI used to create the figure

## Figure Supplements

Figure 4 – Figure Supplement 1: Visualization of SERIA model per participant and linear regressions of behavioral outcomes of the antisaccade task and SERIA predictions

Figure 5 – Figure Supplement 1: Preparatory EEG dynamics in the lateral prefrontal ROI

## Additional Supplementary Files

**Supplementary File 1:** Supplementary Methods: sample size calculation and individual DBS programs for each participant.

**Supplementary File 2:** Supplementary Results of additional trial-by-trial regressions between antisaccade measures and preparatory theta / beta power

## Notes

### Competing Interest Statement

The authors have declared no competing interest.

## REFERENCES

Abbasi, O., Hirschmann, J., Storzer, L., Özkurt, T. E., Elben, S., Vesper, J., Wojtecki, L., Schmitz, G., Schnitzler, A., & Butz, M. (2018). Unilateral deep brain stimulation suppresses alpha and beta oscillations in sensorimotor cortices. NeuroImage, 174. https://doi.org/10.1016/j.neuroimage.2018.03.026

Alegre, M., Lopez-Azcarate, J., Obeso, I., Wilkinson, L., Rodriguez-Oroz, M. C., Valencia, M., Garcia-Garcia, D., Guridi, J., Artieda, J., Jahanshahi, M., & Obeso, J. A. (2013). The subthalamic nucleus is involved in successful inhibition in the stop-signal task: A local field potential study in Parkinson’s disease. Experimental Neurology. https://doi.org/10.1016/j.expneurol.2012.08.027

Allen, D. P. (2009). A frequency domain Hampel filter for blind rejection of sinusoidal interference from electromyograms. Journal of Neuroscience Methods, 177(2). https://doi.org/10.1016/j.jneumeth.2008.10.019

Allen, D. P., Stegemöller, E. L., Zadikoff, C., Rosenow, J. M., & MacKinnon, C. D. (2010). Suppression of deep brain stimulation artifacts from the electroencephalogram by frequency-domain Hampel filtering. Clinical Neurophysiology, 121(8). https://doi.org/10.1016/j.clinph.2010.02.156

Antoniades, C. A., Demeyere, N., Kennard, C., Humphreys, G. W., & Hu, M. T. (2015a). Antisaccades and executive dysfunction in early drug-naive Parkinson’s disease: The discovery study. Movement Disorders, 30(6), 843–847. https://doi.org/10.1002/mds.26134

Antoniades, C. A., Demeyere, N., Kennard, C., Humphreys, G. W., & Hu, M. T. (2015b). Antisaccades and executive dysfunction in early drug-naive Parkinson’s disease: The discovery study. Movement Disorders. https://doi.org/10.1002/mds.26134

Aponte, E. A., Raman, S., Sengupta, B., Penny, W. D., Stephan, K. E., & Heinzle, J. (2016). Mpdcm: A toolbox for massively parallel dynamic causal modeling. Journal of Neuroscience Methods, 257. https://doi.org/10.1016/j.jneumeth.2015.09.009

Aponte, E. A., Schöbi, D., Stephan, K. E., & Heinzle, J. (2017). The Stochastic Early Reaction, Inhibition, and late Action (SERIA) model for antisaccades. PLoS Computational Biology, 13(8). https://doi.org/10.1371/journal.pcbi.1005692

Aponte, E. A., Tschan, D. G., Stephan, K. E., & Heinzle, J. (2018). Inhibition failures and late errors in the antisaccade task: Influence of cue delay. Journal of Neurophysiology, 120(6). https://doi.org/10.1152/jn.00240.2018

Babapoor-Farrokhran, S., Vinck, M., Womelsdorf, T., & Everling, S. (2017). Theta and beta synchrony coordinate frontal eye fields and anterior cingulate cortex during sensorimotor mapping. Nature Communications. https://doi.org/10.1038/ncomms13967

Bakhtiari, S., Altinkaya, A., Pack, C. C., & Sadikot, A. F. (2020). The Role of the Subthalamic Nucleus in Inhibitory Control of Oculomotor Behavior in Parkinson’s Disease. Scientific Reports. https://doi.org/10.1038/s41598-020-61572-4

Ballanger, B., Van Eimeren, T., Moro, E., Lozano, A. M., Hamani, C., Boulinguez, P., Pellecchia, G., Houle, S., Poon, Y. Y., Lang, A. E., & Strafella, A. P. (2009). Stimulation of the subthalamic nucleus and impulsivity: Release your horses. Annals of Neurology, 66(6). https://doi.org/10.1002/ana.21795

Bari, A., & Robbins, T. W. (2013). Inhibition and impulsivity: Behavioral and neural basis of response control. In Progress in Neurobiology. https://doi.org/10.1016/j.pneurobio.2013.06.005

Bates, D. (2005). Fitting linear mixed models in R. Using the lme4 package. R News, 5. https://doi.org/10.1159/000323281

Benis, D., David, O., Lachaux, J. P., Seigneuret, E., Krack, P., Fraix, V., Chabardès, S., & Bastin, J. (2014). Subthalamic nucleus activity dissociates proactive and reactive inhibition in patients with Parkinson’s disease. NeuroImage, 91. https://doi.org/10.1016/j.neuroimage.2013.10.070

Brainard, D. H. (1997). The Psychophysics Toolbox. Spatial Vision, 10(4). https://doi.org/10.1163/156856897X00357

Brown, M. R. G., Vilis, T., & Everling, S. (2007). Frontoparietal activation with preparation for antisaccades. Journal of Neurophysiology. https://doi.org/10.1152/jn.00460.2007

Cavanagh, J. F., & Frank, M. J. (2014). Frontal theta as a mechanism for cognitive control. In Trends in Cognitive Sciences (Vol. 18, Issue 8). https://doi.org/10.1016/j.tics.2014.04.012

Cavanagh, J. F., Wiecki, T. V., Cohen, M. X., Figueroa, C. M., Samanta, J., Sherman, S. J., & Frank, M. J. (2011). Subthalamic nucleus stimulation reverses mediofrontal influence over decision threshold. Nature Neuroscience, 14(11). https://doi.org/10.1038/nn.2925

Chevalier, G., & Deniau, J. M. (1990). Disinhibition as a basic process in the expression of striatal functions. In Trends in Neurosciences (Vol. 13, Issue 7). https://doi.org/10.1016/0166-2236(90)90109-N

Chikazoe, J., Konishi, S., Asari, T., Jimura, K., & Miyashita, Y. (2007). Activation of right inferior frontal gyrus during response inhibition across response modalities. Journal of Cognitive Neuroscience, 19(1). https://doi.org/10.1162/jocn.2007.19.1.69

Cohen, M. X., & Cavanagh, J. F. (2011). Single-trial regression elucidates the role of prefrontal theta oscillations in response conflict. Frontiers in Psychology, 2(FEB). https://doi.org/10.3389/fpsyg.2011.00030

Cohen, M. X., Ridderinkhof, K. R., Haupt, S., Elger, C. E., & Fell, J. (2008). Medial frontal cortex and response conflict: Evidence from human intracranial EEG and medial frontal cortex lesion. Brain Research, 1238. https://doi.org/10.1016/j.brainres.2008.07.114

Combs, H. L., Folley, B. S., Berry, D. T. R., Segerstrom, S. C., Han, D. Y., Anderson-Mooney, A. J., Walls, B. D., & van Horne, C. (2015). Cognition and Depression Following Deep Brain Stimulation of the Subthalamic Nucleus and Globus Pallidus Pars Internus in Parkinson’s Disease: A Meta-Analysis. In Neuropsychology Review (Vol. 25, Issue 4). https://doi.org/10.1007/s11065-015-9302-0

Conway, Z. J., Silburn, P. A., Thevathasan, W., Maley, K. O., Naughton, G. A., & Cole, M. H. (2019). Alternate Subthalamic Nucleus Deep Brain Stimulation Parameters to Manage Motor Symptoms of Parkinson’s Disease: Systematic Review and Meta-analysis. In Movement Disorders Clinical Practice (Vol. 6, Issue 1). https://doi.org/10.1002/mdc3.12681

Cooper, P. S., Karayanidis, F., McKewen, M., McLellan-Hall, S., Wong, A. S. W., Skippen, P., & Cavanagh, J. F. (2019). Frontal theta predicts specific cognitive control-induced behavioural changes beyond general reaction time slowing. NeuroImage, 189. https://doi.org/10.1016/j.neuroimage.2019.01.022

Cordones, I., Gómez, C. M., & Escudero, M. (2013). Cortical Dynamics during the Preparation of Antisaccadic and Prosaccadic Eye Movements in Humans in a Gap Paradigm. PLoS ONE, 8(5). https://doi.org/10.1371/journal.pone.0063751

De Pretto, M., Mouthon, M., Debove, I., Pollo, C., Schüpbach, M., Spierer, L., & Accolla, E. A. (2021). Proactive inhibition is not modified by deep brain stimulation for Parkinson’s disease: An electrical neuroimaging study. Human Brain Mapping. https://doi.org/10.1002/hbm.25530

Delorme, A., Sejnowski, T., & Makeig, S. (2007). Enhanced detection of artifacts in EEG data using higher-order statistics and independent component analysis. NeuroImage, 34(4). https://doi.org/10.1016/j.neuroimage.2006.11.004

Devos, D., Labyt, E., Derambure, P., Bourriez, J. L., Cassim, F., Reyns, N., Blond, S., Guieu, J. D., Destée, A., & Defebvre, L. (2004). Subthalamic nucleus stimulation modulates motor cortex oscillatory activity in Parkinson’s disease. Brain, 127(2). https://doi.org/10.1093/brain/awh053

Emre, M., Aarsland, D., Brown, R., Burn, D. J., Duyckaerts, C., Mizuno, Y., Broe, G. A., Cummings, J., Dickson, D. W., Gauthier, S., Goldman, J., Goetz, C., Korczyn, A., Lees, A., Levy, R., Litvan, I., McKeith, I., Olanow, W., Poewe, W., … Dubois, B. (2007). Clinical diagnostic criteria for dementia associated with Parkinson’s disease. In Movement Disorders (Vol. 22, Issue 12). https://doi.org/10.1002/mds.21507

Engel, A. K., & Fries, P. (2010). Beta-band oscillations-signalling the status quo? In Current Opinion in Neurobiology (Vol. 20, Issue 2). https://doi.org/10.1016/j.conb.2010.02.015

Ewenczyk, C., Mesmoudi, S., Gallea, C., Welter, M. L., Gaymard, B., Demain, A., Yahia Cherif, L., Degos, B., Benali, H., Pouget, P., Poupon, C., Lehericy, S., Rivaud-Péchoux, S., & Vidailhet, M. (2017). Antisaccades in Parkinson disease: A new marker of postural control? Neurology. https://doi.org/10.1212/WNL.0000000000003658

Forstmann, B. U., Dutilh, G., Brown, S., Neumann, J., Von Cramon, D. Y., Ridderinkhof, K. R., & Wagenmakers, E. J. (2008). Striatum and pre-SMA facilitate decision-making under time pressure. Proceedings of the National Academy of Sciences of the United States of America, 105(45). https://doi.org/10.1073/pnas.0805903105

Frank, M. J. (2006). Hold your horses: A dynamic computational role for the subthalamic nucleus in decision making. Neural Networks, 19(8). https://doi.org/10.1016/j.neunet.2006.03.006

Gallea, C., Wicki, B., Ewenczyk, C., Rivaud-Péchoux, S., Yahia-Cherif, L., Pouget, P., Vidailhet, M., & Hainque, E. (2021). Antisaccade, a predictive marker for freezing of gait in Parkinson’s disease and gait/gaze network connectivity. Brain : A Journal of Neurology, 144(2). https://doi.org/10.1093/brain/awaa407

Godefroy, O., Azouvi, P., Robert, P., Roussel, M., Legall, D., & Meulemans, T. (2010). Dysexecutive syndrome: Diagnostic criteria and validation study. Annals of Neurology, 68(6). https://doi.org/10.1002/ana.22117

Goetz, C. G., Fahn, S., Martinez-Martin, P., Poewe, W., Sampaio, C., Stebbins, G. T., Stern, M. B., Tilley, B. C., Dodel, R., Dubois, B., Holloway, R., Jankovic, J., Kulisevsky, J., Lang, A. E., Lees, A., Leurgans, S., LeWitt, P. A., Nyenhuis, D., Olanow, C. W., … LaPelle, N. (2007). Movement disorder society-sponsored revision of the unified Parkinson’s disease rating scale (MDS-UPDRS): Process, format, and clinimetric testing plan. Movement Disorders. https://doi.org/10.1002/mds.21198

Gramfort, A., Luessi, M., Larson, E., Engemann, D. A., Strohmeier, D., Brodbeck, C., Goj, R., Jas, M., Brooks, T., Parkkonen, L., & Hämäläinen, M. (2013). MEG and EEG data analysis with MNE-Python. Frontiers in Neuroscience. https://doi.org/10.3389/fnins.2013.00267

Hamm, J. P., Dyckman, K. A., McDowell, J. E., & Clementz, B. A. (2012). Pre-cue fronto-occipital alpha phase and distributed cortical oscillations predict failures of cognitive control. Journal of Neuroscience, 32(20). https://doi.org/10.1523/JNEUROSCI.5198-11.2012

Hauser, T. U., Iannaccone, R., Stämpfli, P., Drechsler, R., Brandeis, D., Walitza, S., & Brem, S. (2014). The feedback-related negativity (FRN) revisited: New insights into the localization, meaning and network organization. NeuroImage, 84. https://doi.org/10.1016/j.neuroimage.2013.08.028

Hershey, T., Revilla, F. J., Wernle, A., Gibson, P. S., Dowling, J. L., & Perlmutter, J. S. (2004). Stimulation of STN impairs aspects of cognitive control in PD. Neurology, 62(7). https://doi.org/10.1212/01.WNL.0000118202.19098.10

Herz, D. M., Little, S., Pedrosa, D. J., Tinkhauser, G., Cheeran, B., Foltynie, T., Bogacz, R., & Brown, P. (2018). Mechanisms Underlying Decision-Making as Revealed by Deep-Brain Stimulation in Patients with Parkinson’s Disease. Current Biology, 28(8). https://doi.org/10.1016/j.cub.2018.02.057

Hinault, T., Larcher, K., Zazubovits, N., Gotman, J., & Dagher, A. (2019). Spatio–temporal patterns of cognitive control revealed with simultaneous electroencephalography and functional magnetic resonance imaging. Human Brain Mapping. https://doi.org/10.1002/hbm.24356

Hwang, K., Ghuman, A. S., Manoach, D. S., Jones, S. R., & Luna, B. (2014). Cortical neurodynamics of inhibitory control. Journal of Neuroscience. https://doi.org/10.1523/JNEUROSCI.4889-13.2014

Jahanshahi, M., Ardouin, C. M. A., Brown, R. G., Rothwell, J. C., Obeso, J., Albanese, A., Rodriguez-Oroz, M. C., Moro, E., Benabid, A. L., Pollak, P., & Limousin-Dowsey, P. (2000). The impact of deep brain stimulation on executive function in Parkinson’s disease. Brain, 123(6). https://doi.org/10.1093/brain/123.6.1142

Jahanshahi, Marjan, Obeso, I., Baunez, C., Alegre, M., & Krack, P. (2015). Parkinson’s disease, the subthalamic nucleus, inhibition, and impulsivity. In Movement Disorders. https://doi.org/10.1002/mds.26049

Jamadar, S. D., Fielding, J., & Egan, G. F. (2013). Quantitative meta-analysis of fMRI and PET studies reveals consistent activation in fronto-striatal-parietal regions and cerebellum during antisaccades and prosaccades. Frontiers in Psychology. https://doi.org/10.3389/fpsyg.2013.00749

Jenkinson, N., & Brown, P. (2011). New insights into the relationship between dopamine, beta oscillations and motor function. In Trends in Neurosciences. https://doi.org/10.1016/j.tins.2011.09.003

Koss, A. M., Alterman, R. L., Tagliati, M., Shils, J. L., Moro, E., Lang, A. E., Strafella, A. P., Poon, Y. Y. W., Arango, P. M., Dagher, A., Hutchison, W. D., & Lozano, A. M. (2005). Calculating total electrical energy delivered by deep brain stimulation systems [1] (multiple letters). In Annals of Neurology (Vol. 58, Issue 1). https://doi.org/10.1002/ana.20525

Kudlicka, A., Clare, L., & Hindle, J. V. (2011). Executive functions in Parkinson’s disease: Systematic review and meta-analysis. In Movement Disorders (Vol. 26, Issue 13, pp. 2305–2315). https://doi.org/10.1002/mds.23868

Kuznetsova, A., Brockhoff, P. B., & Christensen, R. H. B. (2016). lmerTest: Tests for random and fixed effects for linear mixed effect models. In R package version.

Liebrand, M., Pein, I., Tzvi, E., & Krämer, U. M. (2017). Temporal dynamics of proactive and reactive motor inhibition. Frontiers in Human Neuroscience, 11. https://doi.org/10.3389/fnhum.2017.00204

Maris, E., & Oostenveld, R. (2007). Nonparametric statistical testing of EEG- and MEG-data. Journal of Neuroscience Methods. https://doi.org/10.1016/j.jneumeth.2007.03.024

Marks, W. J. (2015). Deep brain stimulation management. In Deep Brain Stimulation Management. https://doi.org/10.1017/CBO9781316026625

Moon, S. Y., Barton, J. J. S., Mikulski, S., Polli, F. E., Cain, M. S., Vangel, M., Hämäläinen, M. S., & Manoach, D. S. (2007). Where left becomes right: A magnetoencephalographic study of sensorimotor transformation for antisaccades. NeuroImage. https://doi.org/10.1016/j.neuroimage.2007.04.040

Moreau, C., Defebvre, L., Destée, A., Bleuse, S., Clement, F., Blatt, J. L., Krystkowiak, P., & Devos, D. (2008). STN-DBS frequency effects on freezing of gait in advanced Parkinson disease. Neurology, 71(2). https://doi.org/10.1212/01.wnl.0000303972.16279.46

Moro, E., Esselink, R. J. A., Xie, J., Hommel, M., Benabid, A. L., & Pollak, P. (2002). The impact on Parkinson’s disease of electrical parameter settings in STN stimulation. Neurology, 59(5). https://doi.org/10.1212/WNL.59.5.706

Munoz, M. J., Goelz, L. C., Pal, G. D., Karl, J. A., Verhagen Metman, L., Sani, S., Rosenow, J. M., Ciolino, J. D., Kurani, A. S., Corcos, D. M., & David, F. J. (2021). Increased Subthalamic Nucleus Deep Brain Stimulation Amplitude Impairs Inhibitory Control of Eye Movements in Parkinson’s Disease. Neuromodulation. https://doi.org/10.1111/ner.13476

Nasreddine, Z. S., Phillips, N. A., Bédirian, V., Charbonneau, S., Whitehead, V., Collin, I., Cummings, J. L., & Chertkow, H. (2005). The Montreal Cognitive Assessment, MoCA: A brief screening tool for mild cognitive impairment. Journal of the American Geriatrics Society. https://doi.org/10.1111/j.1532-5415.2005.53221.x

Nemanich, S. T., & Earhart, G. M. (2016). Freezing of gait is associated with increased saccade latency and variability in Parkinson’s disease. Clinical Neurophysiology. https://doi.org/10.1016/j.clinph.2016.03.017

Obeso, I., Wilkinson, L., Casabona, E., Bringas, M. L., Álvarez, M., Álvarez, L., Pavón, N., Rodríguez-Oroz, M. C., Macías, R., Obeso, J. A., & Jahanshahi, M. (2011). Deficits in inhibitory control and conflict resolution on cognitive and motor tasks in Parkinson’s disease. Experimental Brain Research. https://doi.org/10.1007/s00221-011-2736-6

Obeso, I., Wilkinson, L., Rodríguez-Oroz, M. C., Obeso, J. A., & Jahanshahi, M. (2013). Bilateral stimulation of the subthalamic nucleus has differential effects on reactive and proactive inhibition and conflict-induced slowing in Parkinson’s disease. Experimental Brain Research, 226(3). https://doi.org/10.1007/s00221-013-3457-9

Oswal, A., Litvak, V., Sauleau, P., & Brown, P. (2012). Beta reactivity, prospective facilitation of executive processing, and its dependence on dopaminergic therapy in Parkinson’s disease. Journal of Neuroscience, 32(29). https://doi.org/10.1523/JNEUROSCI.0275-12.2012

Pa, J., Dutt, S., Mirsky, J. B., Heuer, H. W., Keselman, P., Kong, E., Trujillo, A., Gazzaley, A., Kramer, J. H., Seeley, W. W., Miller, B. L., & Boxer, A. L. (2014). The functional oculomotor network and saccadic cognitive control in healthy elders. NeuroImage. https://doi.org/10.1016/j.neuroimage.2014.03.051

Perrin, F., Pernier, J., Bertrand, O., & Echallier, J. F. (1989). Spherical splines for scalp potential and current density mapping. Electroencephalography and Clinical Neurophysiology, 72(2). https://doi.org/10.1016/0013-4694(89)90180-6

Postuma, R. B., Berg, D., Stern, M., Poewe, W., Olanow, C. W., Oertel, W., Obeso, J., Marek, K., Litvan, I., Lang, A. E., Halliday, G., Goetz, C. G., Gasser, T., Dubois, B., Chan, P., Bloem, B. R., Adler, C. H., & Deuschl, G. (2015). MDS clinical diagnostic criteria for Parkinson’s disease. Mov Disord. https://doi.org/10.1002/mds.26424

Praamstra, P., & Pope, P. (2007). Slow brain potential and oscillatory EEG manifestations of impaired temporal preparation in Parkinson’s disease. Journal of Neurophysiology, 98(5). https://doi.org/10.1152/jn.00224.2007

R Core Team. (2014). R Core Team (2014). R: A language and environment for statistical computing. R Foundation for Statistical Computing, Vienna, Austria. URL Http://Www.R-Project.Org/.

Ray, N. J., Jenkinson, N., Brittain, J., Holland, P., Joint, C., Nandi, D., Bain, P. G., Yousif, N., Green, A., Stein, J. S., & Aziz, T. Z. (2009). The role of the subthalamic nucleus in response inhibition: Evidence from deep brain stimulation for Parkinson’s disease. Neuropsychologia, 47(13). https://doi.org/10.1016/j.neuropsychologia.2009.06.011

Ray, Nicola J., Brittain, J. S., Holland, P., Joundi, R. A., Stein, J. F., Aziz, T. Z., & Jenkinson, N. (2012). The role of the subthalamic nucleus in response inhibition: Evidence from local field potential recordings in the human subthalamic nucleus. NeuroImage, 60(1). https://doi.org/10.1016/j.neuroimage.2011.12.035

Redgrave, P., Prescott, T. J., & Gurney, K. (1999). The basal ganglia: A vertebrate solution to the selection problem? In Neuroscience (Vol. 89, Issue 4). https://doi.org/10.1016/S0306-4522(98)00319-4

Rivaud-Péchoux, S., Vermersch, A. I., Gaymard, B., Ploner, C. J., Bejjani, B. P., Damier, P., Demeret, S., Agid, Y., & Pierrot-Deseilligny, C. (2000). Improvement of memory guided saccades in parkinsonian patients by high frequency subthalamic nucleus stimulation. Journal of Neurology Neurosurgery and Psychiatry. https://doi.org/10.1136/jnnp.68.3.381

Ruppert, M. C., Greuel, A., Freigang, J., Tahmasian, M., Maier, F., Hammes, J., van Eimeren, T., Timmermann, L., Tittgemeyer, M., Drzezga, A., & Eggers, C. (2021). The default mode network and cognition in Parkinson’s disease: A multimodal resting-state network approach. Human Brain Mapping, 42(8), 2623–2641. https://doi.org/10.1002/HBM.25393

Schaum, M., Pinzuti, E., Sebastian, A., Lieb, K., Fries, P., Mobascher, A., Jung, P., Wibral, M., & Tüscher, O. (2021). Right inferior frontal gyrus implements motor inhibitory control via beta-band oscillations in humans. ELife, 10. https://doi.org/10.7554/eLife.61679

Scherrer, S., Smith, A. H., Gowatsky, J., Palmese, C. A., Jimenez-Shahed, J., Kopell, B. H., Mayberg, H. S., & Figee, M. (2020). Impulsivity and Compulsivity After Subthalamic Deep Brain Stimulation for Parkinson’s Disease. In Frontiers in Behavioral Neuroscience (Vol. 14). https://doi.org/10.3389/fnbeh.2020.00047

Singh, A. (2018). Oscillatory activity in the cortico-basal ganglia-thalamic neural circuits in Parkinson’s disease. In European Journal of Neuroscience (Vol. 48, Issue 8). https://doi.org/10.1111/ejn.13853

Singh, A., Richardson, S. P., Narayanan, N., & Cavanagh, J. F. (2018). Mid-frontal theta activity is diminished during cognitive control in Parkinson’s disease. Neuropsychologia, 117. https://doi.org/10.1016/j.neuropsychologia.2018.05.020

Stevens, M. C., Kiehl, K. A., Pearlson, G. D., & Calhoun, V. D. (2007). Functional neural networks underlying response inhibition in adolescents and adults. Behavioural Brain Research, 181(1). https://doi.org/10.1016/j.bbr.2007.03.023

Su, D., Chen, H., Hu, W., Liu, Y., Wang, Z., Wang, X., Liu, G., Ma, H., Zhou, J., & Feng, T. (2018). Frequency-dependent effects of subthalamic deep brain stimulation on motor symptoms in Parkinson’s disease: a meta-analysis of controlled trials. Scientific Reports, 8(1). https://doi.org/10.1038/s41598-018-32161-3

Swann, N., Poizner, H., Houser, M., Gould, S., Greenhouse, I., Cai, W., Strunk, J., George, J., & Aron, A. R. (2011). Deep brain stimulation of the subthalamic nucleus alters the cortical profile of response inhibition in the beta frequency band: A scalp EEG study in parkinson’s disease. Journal of Neuroscience. https://doi.org/10.1523/JNEUROSCI.6135-10.2011

Te Woerd, E. S., Oostenveld, R., Bloem, B. R., De Lange, F. P., & Praamstra, P. (2015). Effects of rhythmic stimulus presentation on oscillatory brain activity: The physiology of cueing in Parkinson’s disease. NeuroImage: Clinical, 9. https://doi.org/10.1016/j.nicl.2015.08.018

Tomlinson, C. L., Stowe, R., Patel, S., Rick, C., Gray, R., & Clarke, C. E. (2010). Systematic review of levodopa dose equivalency reporting in Parkinson’s disease. Movement Disorders. https://doi.org/10.1002/mds.23429

Tsujimoto, T., Shimazu, H., & Isomura, Y. (2006). Direct recording of theta oscillations in primate prefrontal and anterior cingulate cortices. Journal of Neurophysiology, 95(5). https://doi.org/10.1152/jn.00730.2005

Ullsperger, M. (2006). Performance monitoring in neurological and psychiatric patients. International Journal of Psychophysiology, 59(1). https://doi.org/10.1016/j.ijpsycho.2005.06.010

van Driel, J., Swart, J. C., Egner, T., Ridderinkhof, K. R., & Cohen, M. X. (2015). (No) time for control: Frontal theta dynamics reveal the cost of temporally guided conflict anticipation. Cognitive, Affective and Behavioral Neuroscience, 15(4). https://doi.org/10.3758/s13415-015-0367-2

van Noordt, S. J. R., Desjardins, J. A., Gogo, C. E. T., Tekok-Kilic, A., & Segalowitz, S. J. (2017). Cognitive control in the eye of the beholder: Electrocortical theta and alpha modulation during response preparation in a cued saccade task. NeuroImage, 145. https://doi.org/10.1016/j.neuroimage.2016.09.054

van Stockum, S., MacAskill, M., Anderson, T., & Dalrymple-Alford, J. (2008). Don’t look now or look away: Two sources of saccadic disinhibition in Parkinson’s disease? Neuropsychologia. https://doi.org/10.1016/j.neuropsychologia.2008.07.002

Varriale, P., Collomb-Clerc, A., Van Hamme, A., Perrochon, A., Kemoun, G., Sorrentino, G., George, N., Lau, B., Karachi, C., & Welter, M. L. (2018). Decreasing subthalamic deep brain stimulation frequency reverses cognitive interference during gait initiation in Parkinson’s disease. Clinical Neurophysiology, 129(11). https://doi.org/10.1016/j.clinph.2018.07.013

Voon, V., Napier, T. C., Frank, M. J., Sgambato-Faure, V., Grace, A. A., Rodriguez-Oroz, M., Obeso, J., Bezard, E., & Fernagut, P. O. (2017). Impulse control disorders and levodopa-induced dyskinesias in Parkinson’s disease: an update. In The Lancet Neurology. https://doi.org/10.1016/S1474-4422(17)30004-2

Waldthaler, J., Stock, L., Student, J., Sommerkorn, J., Dowiasch, S., & Timmermann, L. (2021). Antisaccades in Parkinson’s Disease: A Meta-Analysis. In Neuropsychology Review. https://doi.org/10.1007/s11065-021-09489-1

Waldthaler, J., Tsitsi, P., & Svenningsson, P. (2019a). Vertical saccades and antisaccades: complementary markers for motor and cognitive impairment in Parkinson’s disease. Npj Parkinson’s Disease, 5(1), 11. https://doi.org/10.1038/s41531-019-0083-7

Waldthaler, J., Tsitsi, P., & Svenningsson, P. (2019b). Vertical saccades and antisaccades: complementary markers for motor and cognitive impairment in Parkinson’s disease. Npj Parkinson’s Disease. https://doi.org/10.1038/s41531-019-0083-7

Walton, C. C., O’Callaghan, C., Hall, J. M., Gilat, M., Mowszowski, L., Naismith, S. L., Burrell, J. R., Shine, J. M., & Lewis, S. J. G. (2015). Antisaccade errors reveal cognitive control deficits in Parkinson’s disease with freezing of gait. Journal of Neurology. https://doi.org/10.1007/s00415-015-7910-5

Wang, C., Ulbert, I., Schomer, D. L., Marinkovic, K., & Halgren, E. (2005). Responses of human anterior cingulate cortex microdomains to error detection, conflict monitoring, stimulus-response mapping, familiarity, and orienting. Journal of Neuroscience, 25(3). https://doi.org/10.1523/JNEUROSCI.4151-04.2005

Weintraub, D., Koester, J., Potenza, M. N., Siderowf, A. D., Stacy, M., Voon, V., Whetteckey, J., Wunderlich, G. R., & Lang, A. E. (2010). Impulse control disorders in Parkinson disease: A cross-sectional study of 3090 patients. Archives of Neurology. https://doi.org/10.1001/archneurol.2010.65

Wiecki, T. V., & Frank, M. J. (2013). A computational model of inhibitory control in frontal cortex and basal ganglia. Psychological Review, 120(2). https://doi.org/10.1037/a0031542

Witt, K., Pulkowski, U., Herzog, J., Lorenz, D., Hamel, W., Deuschl, G., & Krack, P. (2004). Deep Brain Stimulation of the Subthalamic Nucleus Improves Cognitive Flexibility but Impairs Response Inhibition in Parkinson Disease. Archives of Neurology, 61(5). https://doi.org/10.1001/archneur.61.5.697

Wojtecki, L., Timmermann, L., Jörgens, S., Südmeyer, M., Maarouf, M., Treuer, H., Gross, J., Lehrke, R., Koulousakis, A., Voges, J., Sturm, V., & Schnitzler, A. (2006). Frequency-dependent reciprocal modulation of verbal fluency and motor functions in subthalamic deep brain stimulation. Archives of Neurology, 63(9). https://doi.org/10.1001/archneur.63.9.1273

Yugeta, A., Terao, Y., Fukuda, H., Hikosaka, O., Yokochi, F., Okiyama, R., Taniguchi, M., Takahashi, H., Hamada, I., Hanajima, R., & Ugawa, Y. (2010). Effects of STN stimulation on the initiation and inhibition of saccade in parkinson disease. Neurology. https://doi.org/10.1212/WNL.0b013e3181d31e0b

Zaghloul, K. A., Weidemann, C. T., Lega, B. C., Jaggi, J. L., Baltuch, G. H., & Kahana, M. J. (2012). Neuronal activity in the human subthalamic nucleus encodes decision conflict during action selection. Journal of Neuroscience, 32(7). https://doi.org/10.1523/JNEUROSCI.5815-11.2012

Zavala, B. A., Tan, H., Little, S., Ashkan, K., Hariz, M., Foltynie, T., Zrinzo, L., Zaghloul, K. A., & Brown, P. (2014). Midline frontal cortex low-frequency activity drives subthalamic nucleus oscillations during conflict. Journal of Neuroscience, 34(21). https://doi.org/10.1523/JNEUROSCI.1169-14.2014

Zavala, B., Brittain, J. S., Jenkinson, N., Ashkan, K., Foltynie, T., Limousin, P., Zrinzo, L., Green, A. L., Aziz, T., Zaghloul, K., & Brown, P. (2013). Subthalamic nucleus local field potential activity during the eriksen flanker task reveals a novel role for theta phase during conflict monitoring. Journal of Neuroscience, 33(37). https://doi.org/10.1523/JNEUROSCI.1036-13.2013

Zavala, B., Tan, H., Ashkan, K., Foltynie, T., Limousin, P., Zrinzo, L., Zaghloul, K., & Brown, P. (2016). Human subthalamic nucleus-medial frontal cortex theta phase coherence is involved in conflict and error related cortical monitoring. NeuroImage, 137. https://doi.org/10.1016/j.neuroimage.2016.05.031

Zavala, B., Zaghloul, K., & Brown, P. (2015). The subthalamic nucleus, oscillations, and conflict. Movement Disorders, 30(3). https://doi.org/10.1002/mds.26072

Zhang, F., & Iwaki, S. (2019). Common neural network for different functions: An investigation of proactive and reactive inhibition. Frontiers in Behavioral Neuroscience, 13. https://doi.org/10.3389/fnbeh.2019.00124

Zhang, Y., Chen, Y., Bressler, S. L., & Ding, M. (2008). Response preparation and inhibition: The role of the cortical sensorimotor beta rhythm. Neuroscience. https://doi.org/10.1016/j.neuroscience.2008.06.061

